# Understanding and predicting socioeconomic determinants of deforestation in Vietnam’s Central Annamites Landscape (CAL): Pilot study implementing a spatial econometric approach

**DOI:** 10.1101/2021.03.18.436032

**Authors:** Katie P. Bernhard, Stefano Zenobi, Aurélie C. Shapiro

**Affiliations:** United Nations Development Programme, Kampala, Uganda; World Wide Fund for Nature - Greater Mekong, Thailand; Geography Department, Humboldt Universität-zu-Berlin, Berlin, Germany; Forestry Division, Food and Agricultural Organization, Rome, Italy

**Keywords:** Tree cover loss, deforestation, economic development, land conversion, Environmental Kuznets Curve, poverty-environment, spatial econometrics, Vietnam

## Abstract

The forests of the Greater Mekong Subregion, consisting of Myanmar, Thailand, Cambodia, Laos and Vietnam, are under high pressure from economic development and exploitation of natural resources, including but not limited to land concession, smallholder plantations and commercial agriculture, agroforestry development, mining, and road infrastructure development. While these threats are well-known, the magnitude and dynamics of their individual and interacting effects on forest cover are not fully understood. This pilot study aims to apply existing, publicly available macro, micro, and socioeconomic data in addition to remote sensing forest cover data to explore economic determinants of deforestation in the Greater Mekong, using the case of the Central Annamites Landscape (CAL) ecoregion of Vietnam. A longitudinal panel was constructed for 2000-2017, containing 1,658 observations for 144 variables across 95 Tier 2 (district) administrative units in CAL from 2000-2017 for modeling macroeconomic and socioeconomic conditions against annual tree cover loss aggregated to the Tier 2 administrative unit. The first phase of the study used tiered spatial regression analysis to correlatively identify which commodities, economic development activities, and social conditions have historically had the greatest effect on forest cover by magnitude in CAL. Based on first phase results, we selected a subset of these determinants for scenario modelling to predict possible deforestation outcomes given certain economic scenarios. The results, among others, indicate that, when spatially collocated, high poverty rates and smaller scale agricultural land conversion are key immediate determinants of deforestation. This therefore provides evidence to support programs targeting rubber, acacia harvesting, artisanal mining, and land conversion for cash crop plantations. Education is also key immediate socioeconomic factor, as poverty rate is consistently associated with tree cover loss and increased educational attainment is consistently associated with reduced tree cover loss, at approximately 12 ha per percentage increase in secondary school graduation. On the macro level, economic growth in China and Vietnam are correlatively associated with tree cover loss, as are rising trade in Myanmar and Laos. This study provides a methodological contribution to the current academic literature identifying socioeconomic dimensions of deforestation through spatial econometric analysis and scenario modelling at the landscape level. For practitioner work, this pilot provides a model or tool for strategic planning of conservation interventions in light of economic conditions and factors.

## 1. Introduction

The Greater Mekong Subregion, which includes Myanmar, Thailand, Cambodia, Laos and Vietnam, hosts one of the world’s most biodiverse forests while simultaneously supporting a large human population. The region’s forests are under high pressure from economic development and exploitation of natural resources, including but not limited to economic land concession, smallholder plantations and commercial agriculture, agroforestry development, mining, and roads. While these threats are well-known, the magnitude and dynamics of their individual and interacting effects on forest cover are not fully understood.

Between 1990 and 2015, approximately 4.7 million hectares (ha) of forest are reported to have been lost in the Greater Mekong Subregion (GMS), amounting to 2.5% of the total land area (Costenbader et. al, 2015). Average annual decrease in forest cover across the GMS has been approximately 0.21% over the period. Biodiversity loss and species extinctions in the GMS have been attributed to this tree cover loss through habitat fragmentation, degradation, and destruction (WWF-GM Forests Report, 2018). In Vietnam, this tree cover loss has persisted despite reforestation progress in some areas. For example, mountainous regions of Vietnam experienced “forest transition” during the 1990s, with observed overall annual forest cover increases driven largely by reduced forest-to-agriculture conversion as a result of intensification of agriculture on existing land (Meyfroidt and Lambin, 2007; Hansen et al., 2013). However, economic and socioeconomic drivers of deforestation persist, and, as the Central Annamites Landscape (CAL) demonstrates, reforestation is neither spatially nor temporally consistent (Figures 1 and 3). While some subnational regions may undergo this “forest transition,” other regions may still experience deforestation. Indeed, Figure 1 shows average annual tree cover loss in CAL districts from 2001-2017. Reforestation areas may actually accelerate deforestation in other areas by displacement or leakage, when the drivers of deforestation persist (Meyfroidt and Lambin, 2009). Additionally, reforestation may not eliminate forest edge effects and habitat fragmentation, as reforestation does not necessarily ensure density or quality, in terms of biodiversity, of new forest. (Meyfroidt and Lambin, 2008b).

**Figure 1.**
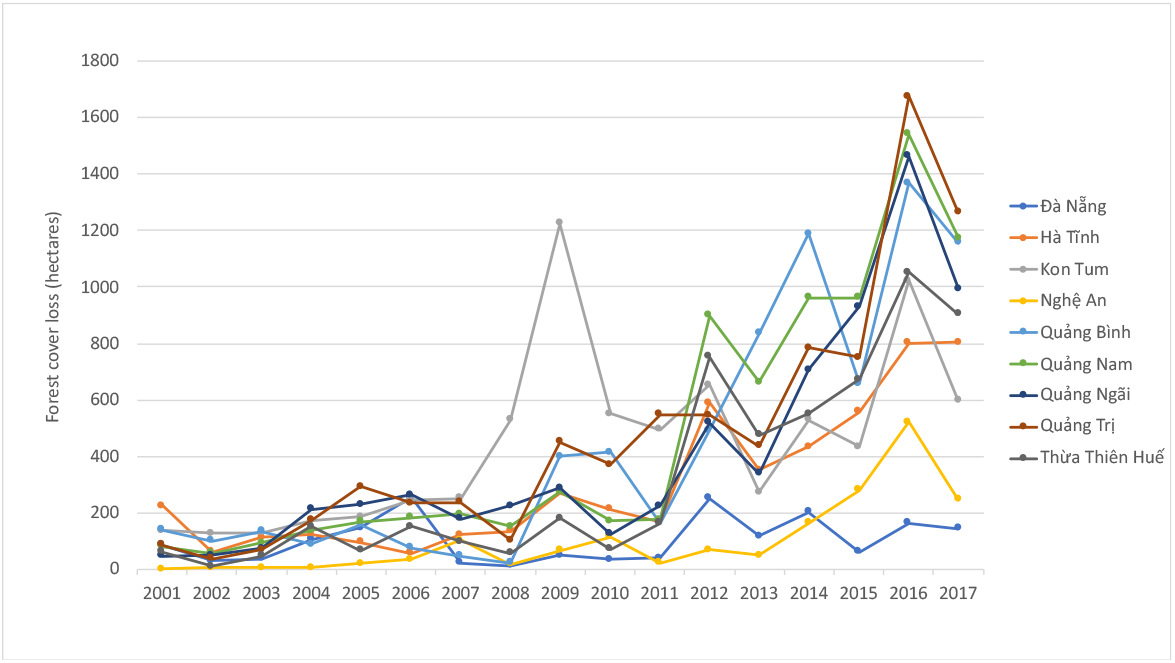
Average tree cover loss trends by CAL province, in hectares, for the landscape area of interest. Overall, tree cover loss increased from 2001-2017 for all provinces, with a slow in tree cover loss 2016-2017 (Data: GFW, 2018).

**Figure 2.**
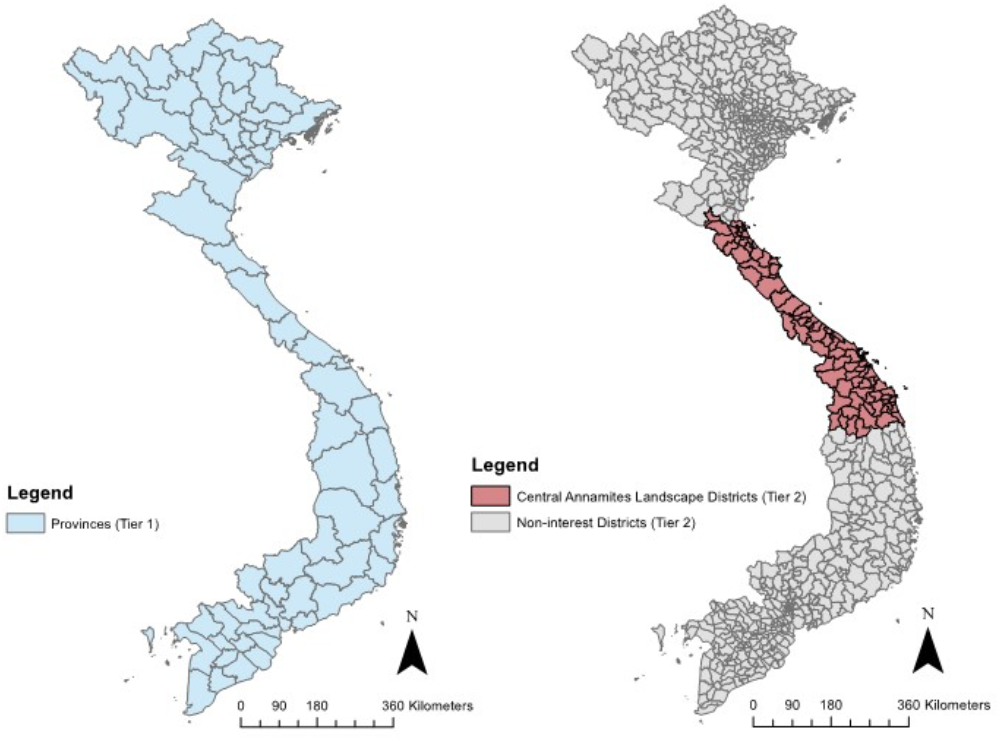
Area of interest in Vietnam. Left: Vietnamese provinces, the Tier 1 administrative level. Right: Vietnamese districts, the Tier 2 administrative level. In red are the Tier 2 districts of interest in the Central Annamites Landscape. These districts are within nine central provinces: Ha Tinh, Nghe An, Quang Binh, Quang Tri, Thua Thien Hue, Da Nang, Quang Nam, Quang Ngai, Kon Tum, and Binh Dinh (Data: GADM, 2016).Binh Dinh was ultimately removed from analysis it has insufficient complete forest cover.

**Figure 3.**
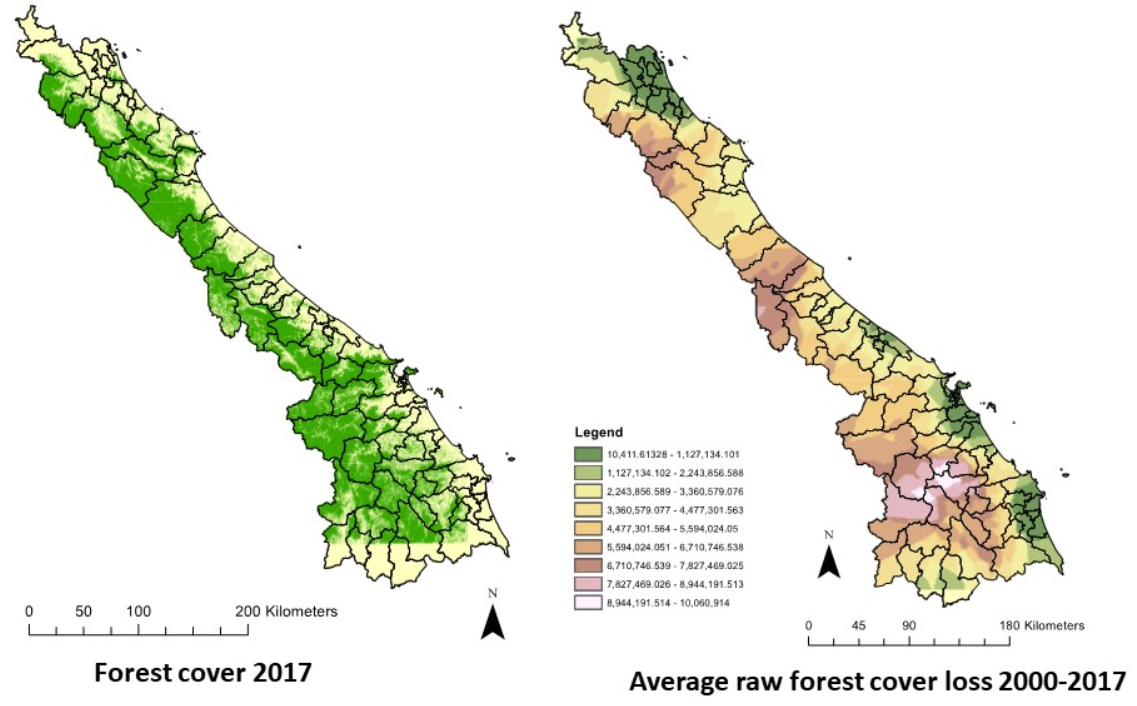
Left: Forest cover in 2017 in the CAL area of interest. Green indicates forest cover. Data: WWF Forest Cover (Data: GFW, 2018). Forest cover and forest cover loss aggregated to district for analysis to account for economic and socioeconomic data. 1m cell. Right: Average annual forest cover loss in the CAL area of interest (sq m), using inverse distance weight (IDW) interpolation of district level forest cover loss averages (power = 0.2 to smooth across districts; interpolated to 5 surrounding points) (Data: GFW, 2018).

The objectives of this pilot study are twofold: First, we sought to systematically assess available macro and microeconomic data that can be used for modeling economic determinants of deforestation in the Central Annamites Landscape (CAL) ecoregion of Vietnam (Figures 2 and 3).

Second, we use these data to conduct initial modeling of specific socioeconomic determinants of deforestation in the CAL, focusing on economic activities such as production of key commodities in addition to social conditions. Our main research questions were the following:

- Which commodities, economic development activities, and social conditions have historically had the greatest effect on forest cover by magnitude in the CAL?
- How do different commodities interact in leading to this deforestation?
- If the current trends of economic growth are maintained, to what extent will deforestation be affected? Spatially, which areas in the CAL are most “at risk”?

To answer these questions, the initial phase of this study identified principal determinants and their historical magnitude and interaction from 2000-2017. Based on the results of the first phase, we selected a subset of these determinants of interest for scenario modelling in order to predict possible deforestation outcomes given scenarios in these interest areas. This study provides a methodological contribution to the current academic literature identifying socioeconomic dimensions of deforestation through spatial econometric analysis and scenario modelling at the landscape level. From the practitioner perspective, we hope that this pilot study provides a useful tool for strategic planning of conservation interventions in light of economic conditions and factors, not only through the empirical analyses conducted but also through our systematic assessment of currently available data for understanding these issues. As a pilot, we hope to highlight areas of need for additional data collection and future research. Lastly, we hope to move beyond or provide empirical justification for the “deforesting commodity of the year” effect for certain commodities based on visible investments and media attention, as this focus does not allow for long term programmatic planning.

### 1.1 Conceptual Framework

There is a robust literature seeking to understand both the drivers and the consequences of deforestation. Similar to several studies, such as Khuc et al. (2018), we utilized the Angelesen and Kaimowitz (1999) theoretical model of socioeconomic determinants of deforestation, further detailed in Figure 4. We selected framework because even alternative models in the literature have in many cases been variations of the Angelsen and Kaimowitz approach, which is a foundational model in the study of socioeconomic determinants of deforestation (Chomitz, 2006). The Angelsen and Kaimowitz framework identifies three levels of determinants: underlying macroeconomic causes, immediate decision parameters, and agent-level decision-making.

**Figure 4.**
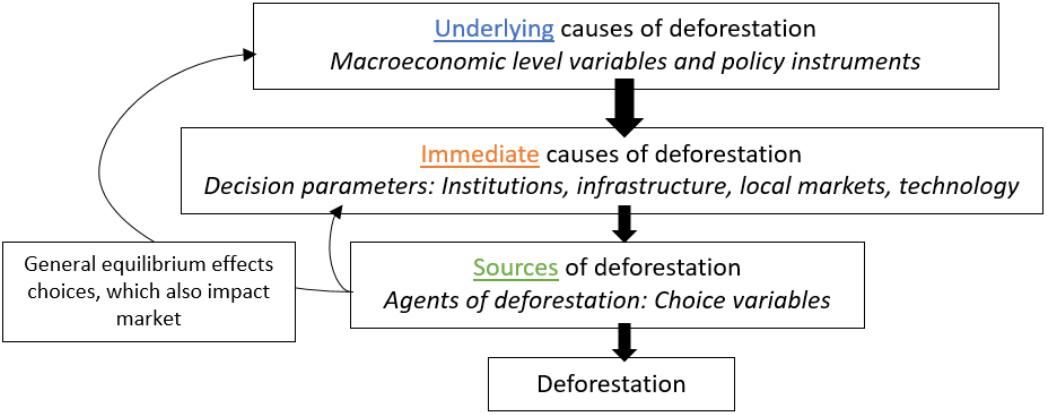
Variables affecting deforestation. Adapted from Angelsen and Kaimowitz (1999).

As we apply this conceptual and model framework in this study, deforestation at location *i* and time *t* is thus a function of underlying and immediate determinants and agent decision parameters,

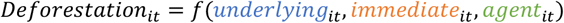

and each of these determinants is, in turn, a nested function of key economic and socioeconomic conditions including commodity prices, GDP per capita, GDP growth, trade liberalization, income and poverty, population density, education levels, off-farm wage opportunities, agricultural suitability of land, and an agent’s background, preferences, and resources.

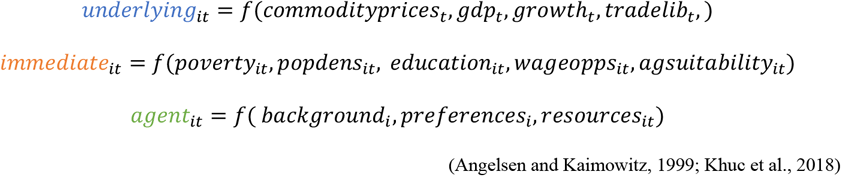

As discussed in Scrieciu (2007), the inclusion of all three levels of determinants in a single model can result in endogeneity and lack of clarity in identifying causal links, as ultimately, variables determining agent decision-making are affected by immediate determinants, in turn effected by underlying determinants. We apply this in our application of the framework and have discussed the variables used in further detail in Table 1. We also review the literature describing our main variables below.

**Table 1.**
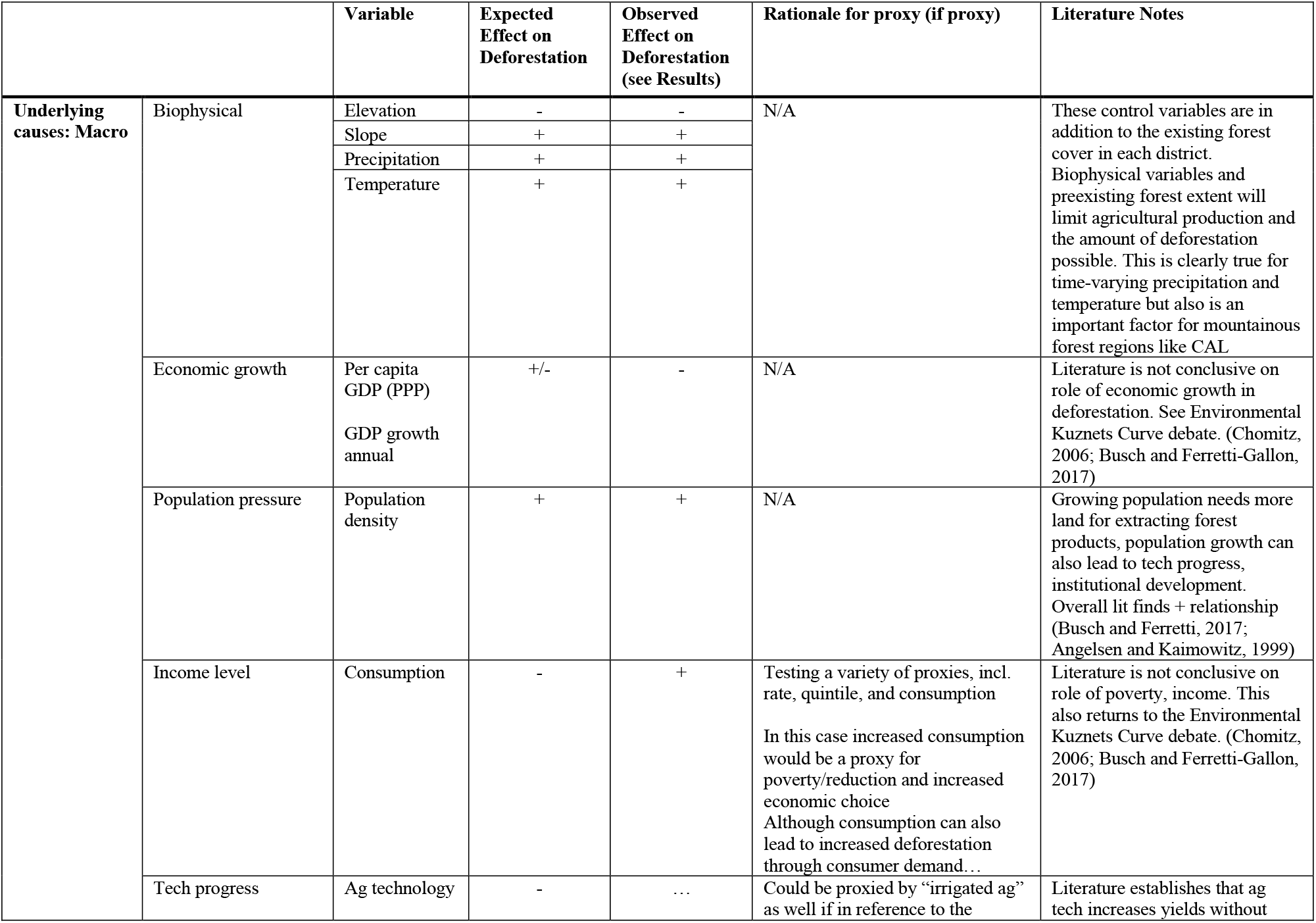
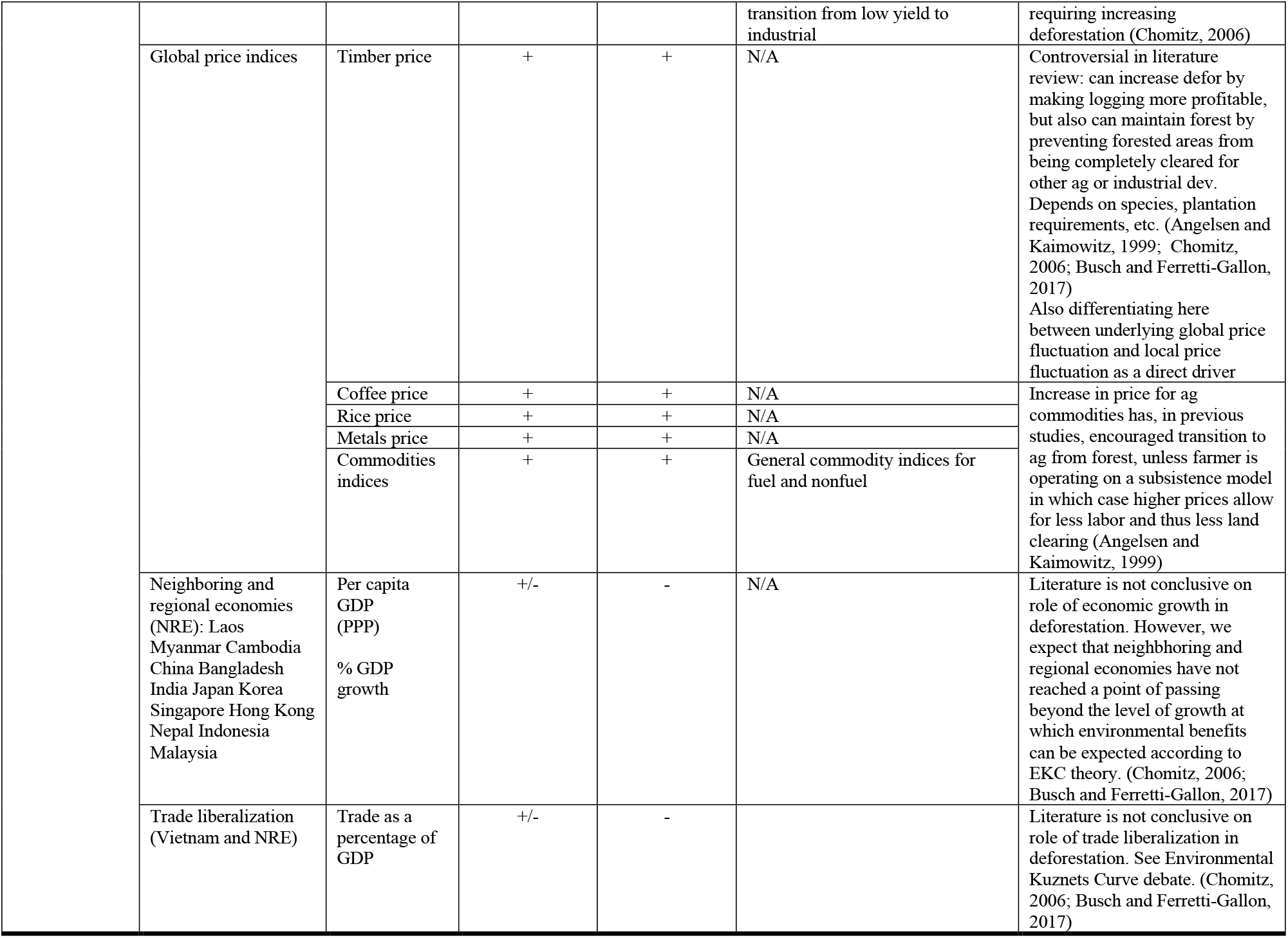
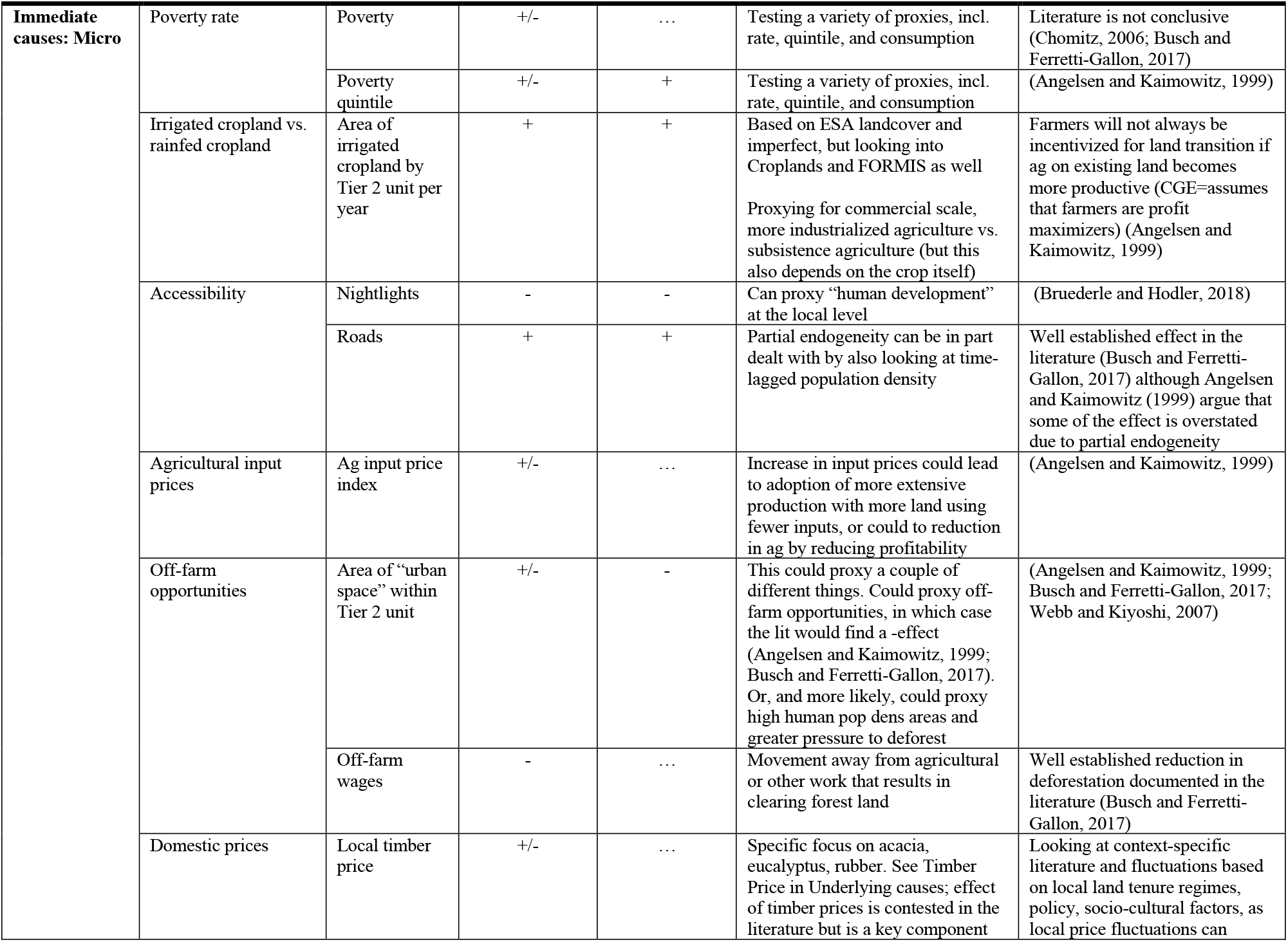
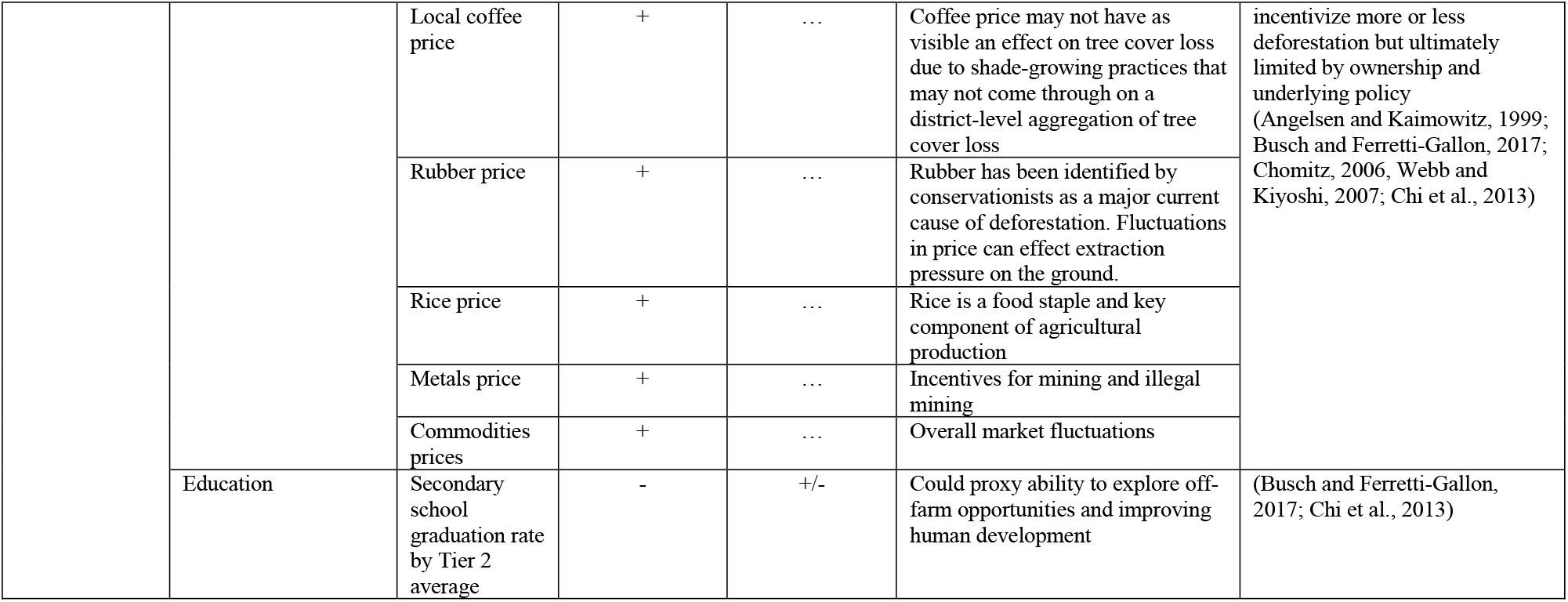
Expected and observed outcomes for driver variables based on CAL context-specific and metanalysis evidence from the landscape change and deforestation literatures. See Summary Statistics (Appendix I) for variable detail and units.

First, poverty-environment linkages have long been the subject of scholarly debate (Chomitz, 2006). Barbier (2017) finds that the relationship between income and poverty and deforestation is complex and often context-dependent. Off-farm employment opportunities are an important determinant for the effect that poverty may have on deforestation (Busch and Ferretti-Gallon, 2017). In Vietnam, the provincial level averages of poverty rates (Figure 5) demonstrate that the overall trend is improving (GSOV, 2018). This decline is the result of several economic and policy mechanisms. In this study we seek to understand both the effect of poverty on deforestation in CAL but also to describe the relationships between some of these economic mechanisms. In addition to provincial-level poverty data, we use the intensity of nighttime lights at the district level to proxy human development and industrial development, as this variable is collinear with road infrastructure. We also proxy access to off-farm wages with the proportion of each district that is covered by urban area.

**Figure 5.**
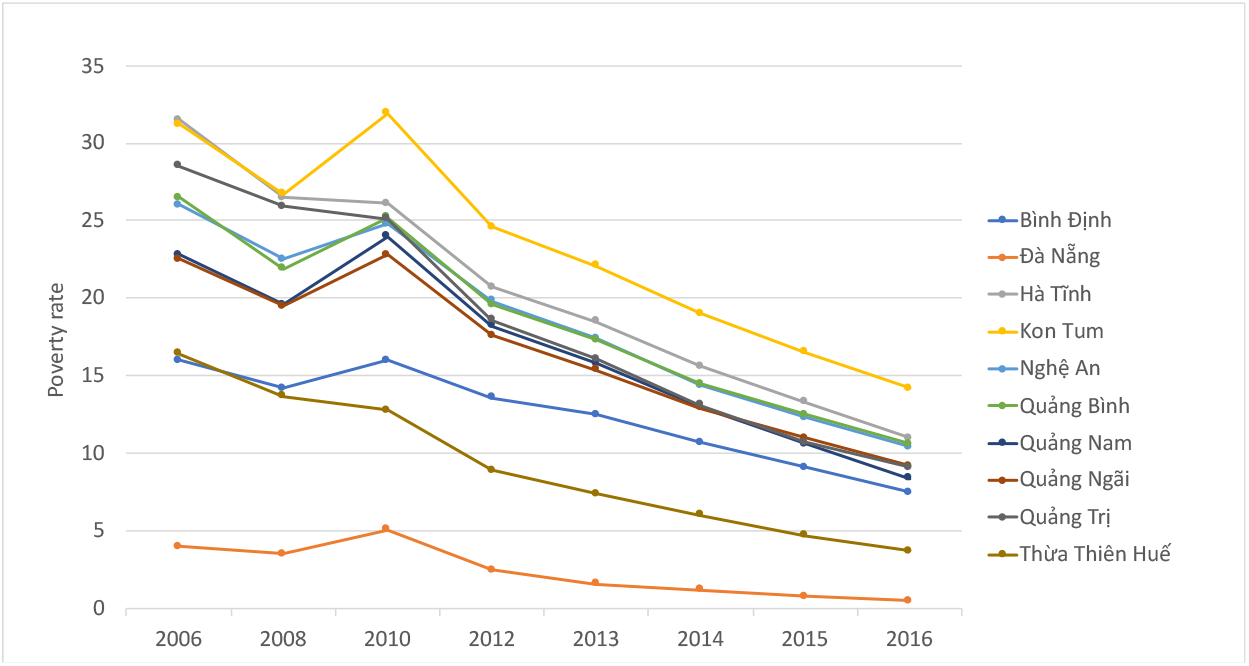
Poverty rate trends by CAL province, according to GSOV data, for roughly 2006-2016 with dropout years. Importantly, GSOV states: “Monthly average income per capita of household which is updated by CPI as follows: 400 thousand dongs for rural areas and 500 thousand dongs for urban areas in 2010; 570 thousand dongs and 710 thousand dongs in 2013; 605 thousand dongs and 750 thousand dongs in 2014; 615 thousand dongs and 760 thousand dongs in 2015 and 630 thousand dongs and 780 thousand dongs in 2016, respectively.” (Data: GSOV)

Similar to poverty-environment linkages, the effect of economic growth and GDP on the environment is contested in the literature. According to Environmental Kuznets Curve (EKC) theories, economic growth degrades the environment until a certain point, after which the benefits of economic growth and development, such as enhanced technology coupled with stronger regulations and institutions, leads to improved environmental outcomes (Panayotou, 1994). However, evidence for this relationship exists only for certain types of environmental degradation, such as air pollution, while the effect on, other degradation types, such as deforestation, is unclear (Cropper and Griffiths, 1994). Bhattari and Hammig (2001) find evidence of a “deforestation EKC” mostly reliant on shifting energy use away from forest-based resources, and improved institutional structures with increased economic development. However, economic growth in Vietnam and its neighbors may have resulted in increasing pressure on forests to support both economic activities and growing populations (Figure 6). Identifying the links between deforestation and economic growth is thus a priority for conservation and the environmental sustainability of economic activities moving forward. In part, this study fits into the EKC literature by assessing the relationship between various indicators of economic growth and development at the landscape level and deforestation.

**Figure 6.**
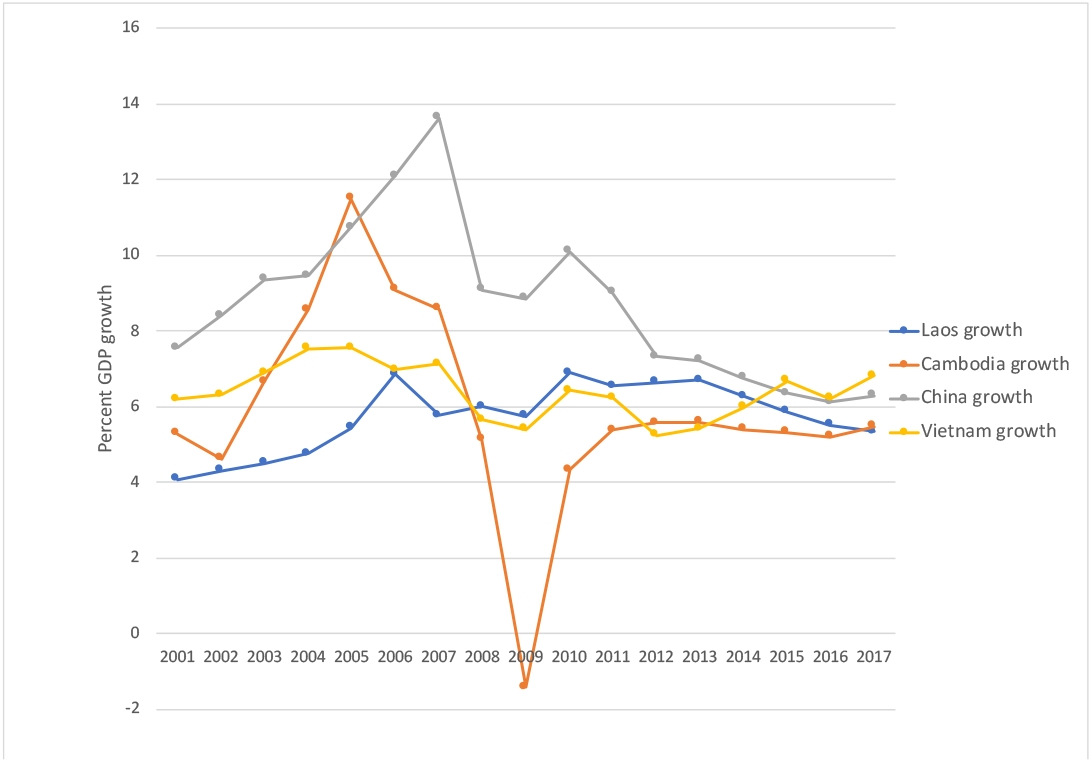
Economic growth from 2001-2017 for Vietnam and selected neighboring economies, Laos, Cambodia, and China. The causes and consequences of negative Cambodian growth rate in 2009 for environmental degradation could also be an interesting focus of future research (Data: WDI, 2018).

Next, increasing human population, particularly increasing population density in proximity to forests and protected areas can be an important determinant of deforestation. Supporting a large human population can lead to more extraction of forest resources, while population density in close proximity to forests, especially in high-poverty areas, can result in community forest dependence (Cropper and Griffiths, 1994; DeFries et al., 2010). Additionally, Pfaff (1999) examined satellite & socio-economic municipality-level data and found deforestation driven by land characteristics, such as soil quality and vegetation type, and factors affecting transport costs such as distance to markets and roads. According to Pfaff, the effects of population growth on deforestation, modeled through migration and credit availability, depended on the preexisting population. We use population density estimates at the district level to understand the dynamics of this population and deforestation in CAL.

Lastly, land use change for agricultural and industrial production has indeed proven to be a key driver of deforestation in CAL. A number of studies have previously used spatial and econometric methods to understand the effect of land use change or socioeconomic conditions on deforestation. These have primarily been at one of two scales: finer, sub-district level, and national-scale. First, several studies have focused on a fine spatial scale, within a single district or province, relying on small scale surveys and/or data collection at the household level (Müller and Zeller, 2002; Webb and Kiyoshi, 2007; Chi et al., 2013). These have not generally incorporated multiple temporal waves due to reliance on limited spatial land use data availability. Müller and Zeller (2002) used spatial econometric methods, supplemented by interviews, to identity drivers of deforestation in the Dak Lak Province specifically in 1975, 1992, and 2000. This study found that deforestation in this province has historically been driven by land-intensive agricultural production. However, as previously discussed, implementing protected areas and “policies discouraging shifting cultivation” in the later 1990s reduced forest pressure while promoting economic growth. (Müller and Zeller, 2002; Meyfroidt and Lambin, 2007). In this study, we hope to contribute insights from CAL from the 2000s.

Second, a number of studies have been conducted at the national scale, incorporating all regions and ecoregions of Vietnam. Khuc et al. (2018) found through spatial and regression-based methods that, at the national level, primary underlying indicators of deforestation and forest degradation include initial forest cover, per capita income, agricultural production, governance, population growth, food, and poverty. Further, the study identified north central, northeast, central highland, and northwest areas of Vietnam as particular risk areas. We use these findings as key support for our identification strategy for econometric analysis in this study.

So far no other studies have sought to quantify the relationship between socioeconomic conditions and deforestation in the CAL at the landscape level using spatially explicit methods. Because the spatially explicit previous studies have also relied on one, or in some cases two, maps of land use change, our study is among the first to use dense time series commodity price data for underlying conditions and as a possible signal of land use change leading to deforestation in Vietnam (Munroe and Müller, 2007). Cropper and Griffiths (1994) found hard log price to be correlated with deforestation at the continent level in Latin America, but neither Africa nor Asia. Scrieciu (2007) found that global commodity price increases are linked to increasing deforestation, but this study was conducted on a broad scale, similar to that of Cropper and Griffiths, using a panel of fifty tropical countries over a period of eighteen years. Figure 7 shows trends in selected commodity prices relative to tree cover loss trends.

**Figure 7.**
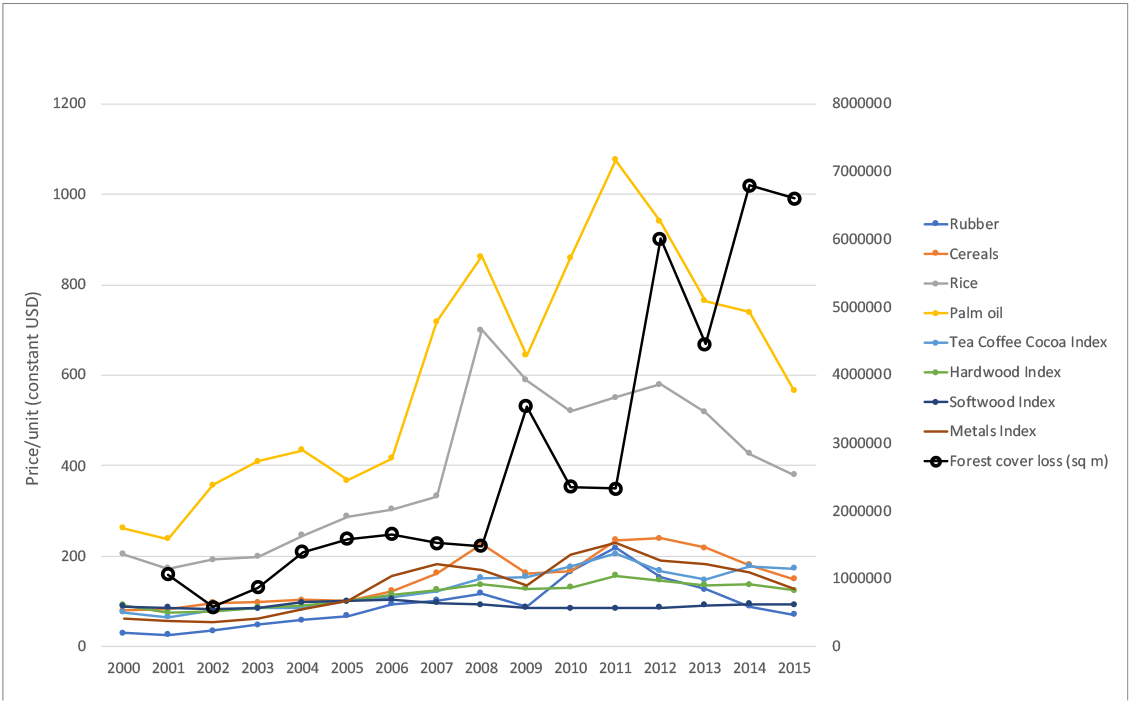
Combination plot, commodity price trends (left axis) relative to tree cover loss in CAL by m^2^ (right axis) over time from 2000-2015 with 2016 and 2017 not included due to dropout (Data: IMF, 2018; GFW, 2018).

In summary, in this study we apply the Angelsen and Kaimowitz (1999) framework for underlying and immediate determinants of deforestation and base our identification and econometric analysis methods on the results of previous spatial econometric studies in Vietnam, such as Khuc et al. (2018). Based on this literature, income and poverty, economic growth, population density, and land use change are key variables in our analysis. Finally, as previously mentioned, Table 1 describes the rationale behind the use of data for specific variables in this study to capture the key socioeconomic conditions emphasized in the literature, and to capture parameters of interest in alignment with the conceptual framework.

## 2. Methods

### 2.1 Data

We constructed a panel of ecological, economic, and social variables for analysis. Prior to constructing this panel, as a key component of this study we systematically reviewed open-source secondary data available for forest cover and economic variables, the findings of which are available in Appendix II. Data availability and quality confined the scope of our study, but also identified important areas for future data collection and research.

The primary response variable used in this study was annual tree cover loss aggregated to the Tier 2 administrative unit. This was extracted from annual raster data acquired from WWF and Global Forest Watch (Hansen, 2013), then aggregated to coarse district-level means. Economic data was sourced from a variety of institutions including IMF, World Bank DataBank, World Database on Protected Areas, the AidData Geoquery platform and University of Delaware, and the General Statistics Office of Vietnam (GSOV). Biophysical factors were included in spatial regression, such as elevation, slope, temperature, and precipitation (aggregated to Tier 2 administrative unit by mean), but time-invariant biophysical factors were not included in site-demeaned fixed effects as those specifications account for time-invariant factors that vary between sites. Proportion of a Tier 2 unit in a protected area, in addition to previously existing forest cover extent as a control variable, were also included in spatial regression. Aggregation to administrative unit rather than grid cells was selected to maintain consistency with the socioeconomic variables from household surveys, which are weighting and representative according to administrative units.

Coarse-scale variables for underlying macroeconomic conditions driving deforestation included per capita GDP (PPP, international), trade as a percentage of GDP, and annual GDP growth for Vietnam and surrounding economies; global commodity prices for commodities relevant to Vietnamese and CAL economies (including but not limited to rubber, rice, coffee, hardwood price indices, softwood price indices, metals price indices, cereals price indices, etc.). We also included annual quantity of foreign direct investment flow delivered to specific districts and earmarked by World Bank. These variables generally exhibited temporal but not spatial variation. For the immediate determinants of deforestation, we include variables which had both spatial and temporal variation, such as population density, poverty rates, proportion of households with access to electricity, district-level contributions to national GDP, proportion of urban area per district, land area devoted to irrigated agriculture per district, nighttime lights, and accessibility by roads. Table 1 outlines the key variables used and expected relationships with deforestation based on the literature and the CAL context.

The panel was constructed for the time period 2000-2017, and thus contains 1,658 observations for 144 variables across 95 Tier 2 (district) administrative units in the CAL from 2000-2017 (Figure 1; Summary Statistics, Appendix I). The CAL boundaries and districts used for analysis were delineated using boundaries provided by WWF-Vietnam. The Tier 2 administrative unit was selected as this provided enough site units within a panel for statistical purposes, but sufficient socioeconomic data was still available for this scale. The panel is unbalanced, as some variables are missing for certain districts or time periods. Nevertheless, the use of panel data enables analysis of temporal and spatial trends in addition to application of a variety of econometric techniques. We first performed simple correlative statistics and exploratory spatial data analysis, which was followed by spatial regression analysis which explored spatially autoregressive and spatial lag of x (SLX) models. Lastly, we utilized the panel nature of the dataset to perform site fixed effects modeling.

### 2.2 Spatial and econometric analysis

We first assessed district-level associations and spatial autocorrelation, particularly for the purposes of constructing spatial lags of key variables exhibiting spatial clustering or dispersion. Spatial autocorrelation was assessed using the univariate local Moran’s I statistic in addition to bivariate local Moran’s I for roughly establishing spatial association between socioeconomic variables and tree cover loss. We chose a row-normalized queen contiguity matrix,

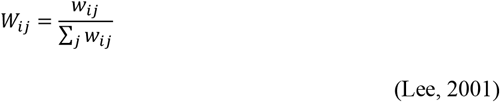

where *w_ij_*=1 indicates contiguity between districts *i* and *j*. Clustering or dispersion is thus assessed using the average of a variable’s values for districts directly bordering the district of interest. As a part of this pilot study, we experimented with a number of alternative spatial weighting options and thus in some instances also utilized a k-nearest neighbor (KNN) approach,

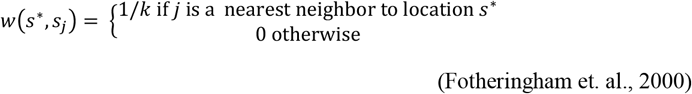

where *w*(*s*\*, *s_j_*), like *W_ij_* above, is a scalar weight assigned to administrative unit *j* based on its distance from location *s*^*^. For KNN weights, a number of “nearest neighbors” is selected to create the weight, essentially a moving average of “neighbor” values for the district of interest. The primary difference to note between the contiguity and nearest neighbor weights is illustrated in Figure 4. In a contiguity weight, the value of a variable at district E would not be included in the lag for district C, even though they are spatially near each other, because they do not share a border. However, in a KNN weight, E would be included in the lag for district C because E is among C’s five nearest neighbors. In CAL, the most common range of frequencies of contiguous neighbors for a given Tier 2 district is 5-7, so we selected five neighbors as the number of averaged neighbors to establish a given district’s weight. We do not use distance-band weights such as Euclidean or inverse distance weighting due to the aggregation of our dataset to administrative units (Anselin, 2018).

**Figure 8.**
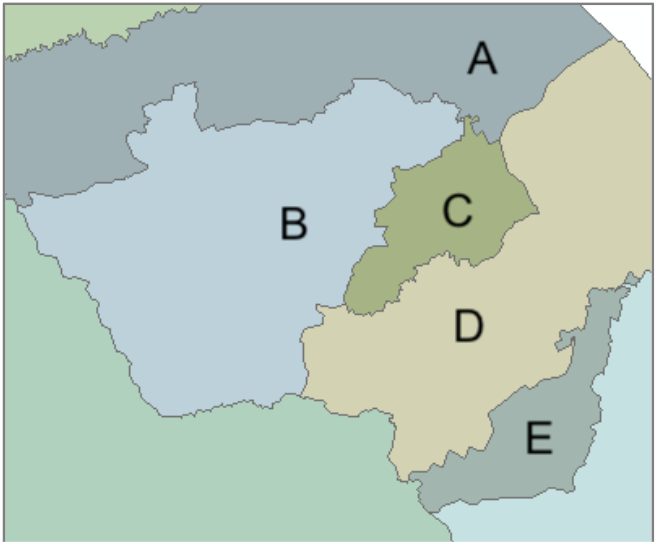
The primary relevant difference between weights structures is the inclusion of non-contiguous near neighbors in the KNN weight such as district E in district C’s lag, unlike the contiguity weight, which would include only neighbor values of A, B, and D in the lag for C.

On a −1 to 1 scale, the univariate Moran’s I statistic compares a variable to its own spatial lag (see Appendix III). The bivariate version of the Moran’s I is mathematically equivalent to the spatial autoregression of *y(Wy)* on *x*, where the relationship between a non-lagged variable and the “neighborhood” of a lagged variable is assessed:

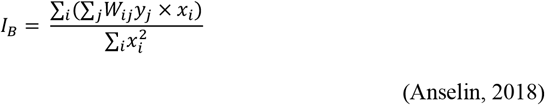

Following the identification of spatial autocorrelation and bivariate spatial association, we then used correlative statistics, including Pearson’s correlation matrices and scatter plot matrices with locally weighted scatterplot smoothing (LOWESS), to better understand the relationships and collinearity between variables used (Table 3; Figures 9 and 10). Next, we performed spatial regression analysis on two separate scales, following the Angelsen and Kaimowitz (1999) approach: for the underlying macroeconomic conditions driving deforestation, and the immediate microeconomic determinants. We separated these two sets of conditions as described by Angelsen and Kaimowitz (1999) and recommended by Scrieciu (2007). As describes in the Literature Review and Figure 4, underlying causes and immediate causes are often endogenous or collinear, as many underlying macroeconomic conditions will ultimately create the conditions for the immediate ones.

**Figure 9.**
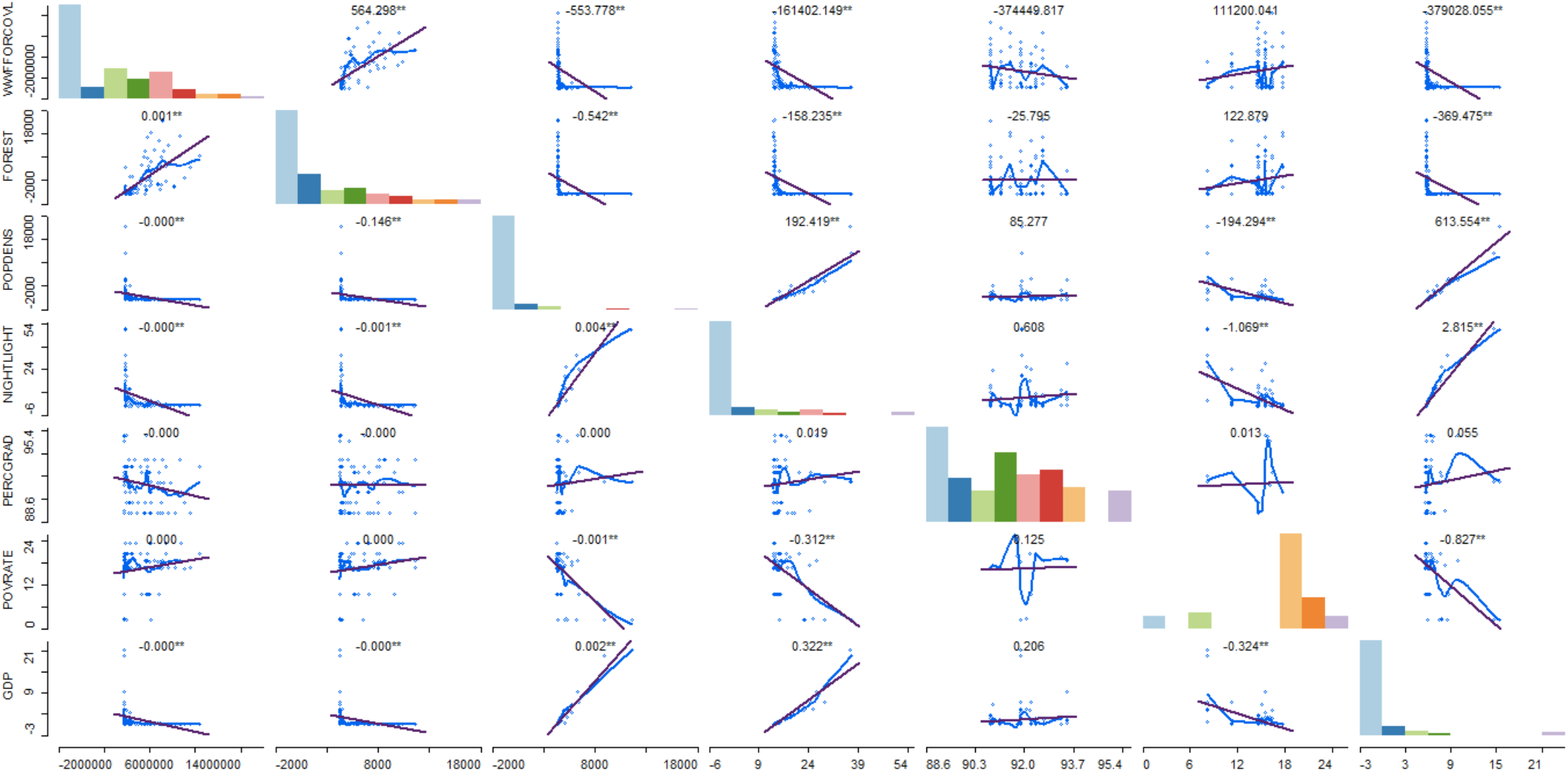
Scatterplot matrices for selected immediate determinants of deforestation variables. Variables on the vertical axes (left-hand side) are the Y-axis variables for each row. The purple line on each scatterplot shows a linear correlation fit, the magnitude and significance of the slop is the number above each plot (e.g., 564.298**). The blue line is a locally weighted scatterplot smoothing (LOWESS) which illustrates the nonlinearities in the relationships between each variable, which can also explain the <0.5 R^2^ values in later regressions. Variables along the bottom are the x-axis variables for each column. Histograms are the distribution of each variable, which can also be helpful for determining which variables require skew-correction to obtain a roughly normal distribution to satisfy Gauss-Markov assumptions for OLS regression. The top row of correlations thus shows the bivariate correlations between forest cover loss and existing forest cover, population density, nighttime lights, secondary school graduation rate, poverty rate, and district contribution to GDP.

**Figure 10.**
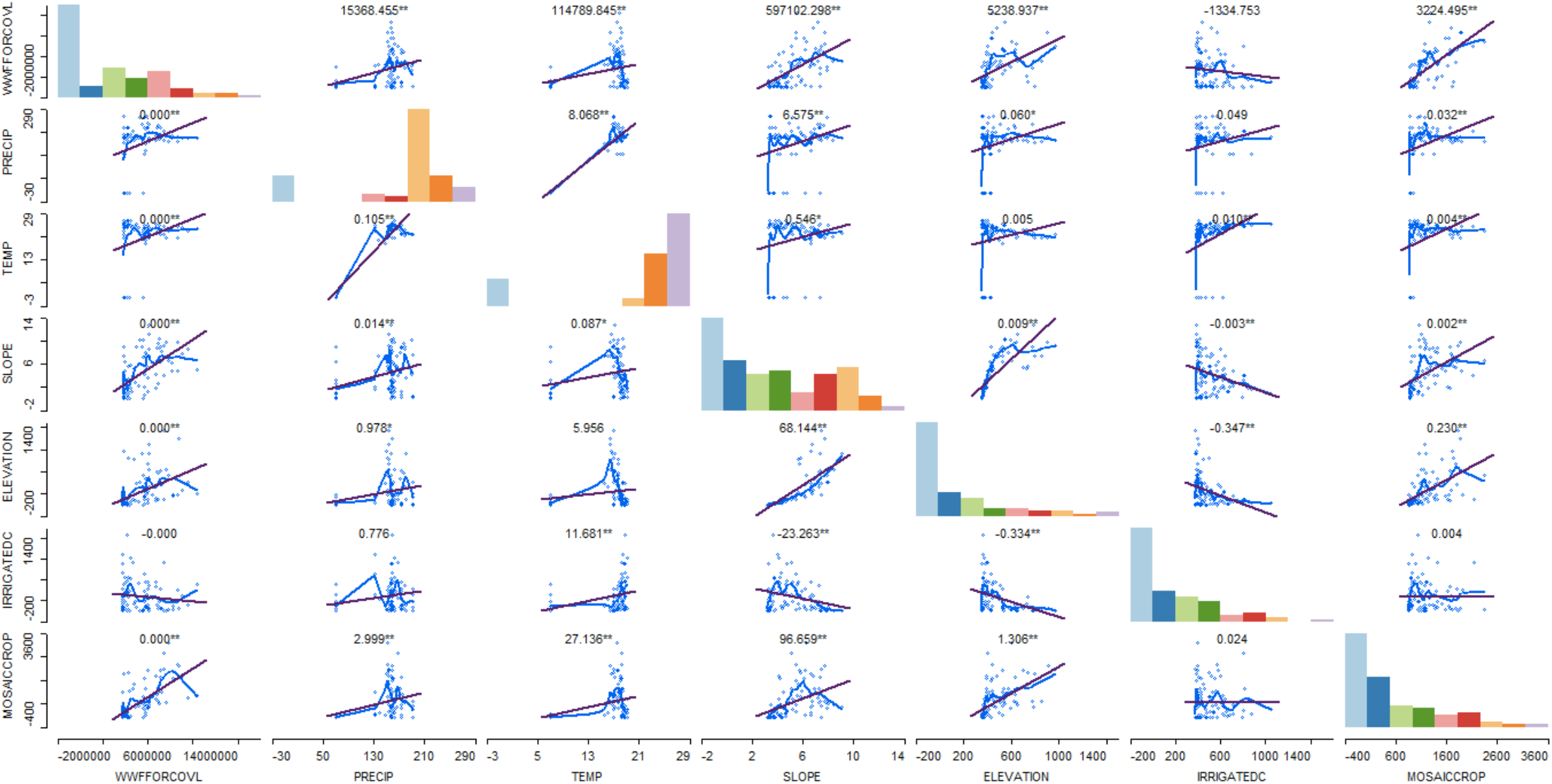
Scatterplot matrices for selected biophysical constraints on deforestation. Variables on the vertical axes (left-hand side) are the Y-axis variables for each row. The purple line on each scatterplot shows a linear correlation fit, the magnitude and significance of the slop is the number above each plot (e.g., 15368.455**). The blue line is a locally weighted scatterplot smoothing (LOWESS) which illustrates the nonlinearities in the relationships between each variable, which can also explain the <0.5 R^2^ values in later regressions. Variables along the bottom are the x-axis variables for each column. Histograms are the distribution of each variable, which can also be helpful for determining which variables require skew-correction to obtain a roughly normal distribution to satisfy Gauss-Markov assumptions for OLS regression. The top row of correlations thus shows the bivariate correlations between forest cover loss and precipitation, temperature, slope, elevation, irrigated cropland, and mosaic cropland. Comparison of the irrigated cropland and mosaic cropland shows no association between forest cover loss and irrigated cropland, but a significant correlation between forest cover loss and mosaic cropland.

**Table 2.**
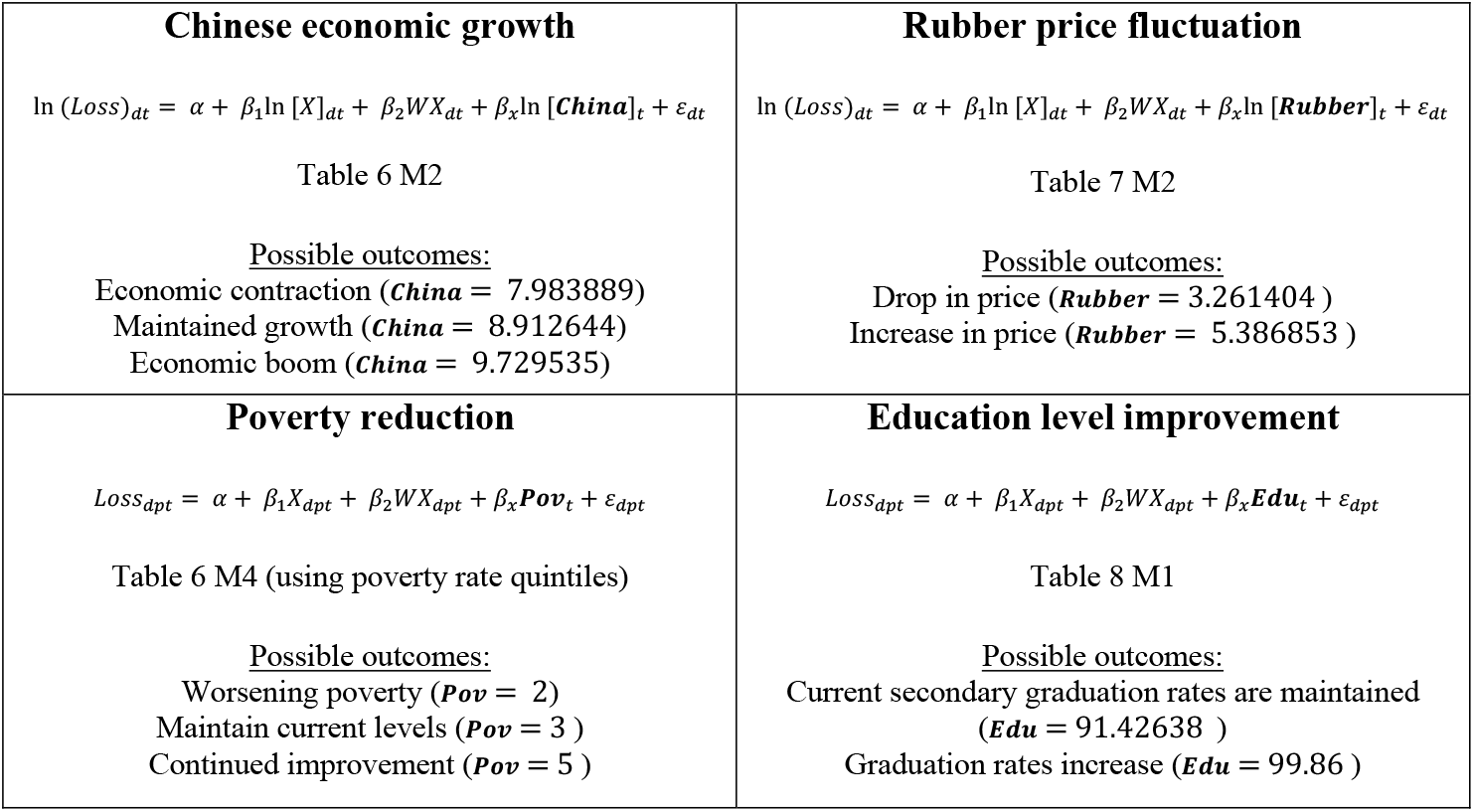
Scenarios fit for Phase 2 using predictive margins based on results of Phase 1 modeling.

**Table 3.** Pearson’s correlation coefficients between Vietnamese GDP and commodity prices/price indices. Values in red indicate correlated variables that are subject to collinearity and are therefore separated in models presented. This due to use of coarse temporal scale and global nature of prices. *** p<0.01, **p<0.05, *p<0.1.

|  | Vietnam GDP | Rice price | Hardwood price index | Softwood price index | Metals price index | Rubber price | Tea coffee cocoa price index | Cereals price index |
| --- | --- | --- | --- | --- | --- | --- | --- | --- |
| Vietnam GDP | 1.000 |  |  |  |  |  |  |  |
| Rice price | 0.6939*** | 1.000 |  |  |  |  |  |  |
| Hardwood price index | 0.8399*** | 0.8873*** | 1.000 |  |  |  |  |  |
| Softwood price index | 0.0358 | -0.1402*** | 0.0017 | 1.000 |  |  |  |  |
| Metals price index | 0.7424*** | 0.8071*** | 0.9499*** | 0.0553** | 1.000 |  |  |  |
| Rubber price | 0.4606*** | 0.7745*** | 0.8805*** | -0.1457*** | 0.9336*** | 1.000 |  |  |
| Tea coffee cocoa price index | 0.8975*** | 0.8359*** | 0.9418*** | -0.1468*** | 0.8699*** | 0.8348*** | 1.000 |  |
| Cereals price index | 0.7707*** | 0.9114*** | 0.9401*** | -0.1380*** | 0.8877*** | 0.8578*** | 0.8616*** | 1.000 |

We modeled underlying macroeconomic determinants of deforestation using the following simple linear OLS estimation with spatially lagged explanatory variables:

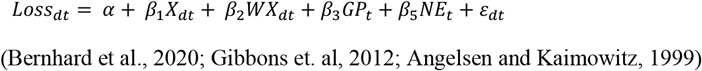

where *Loss_dt_* is tree cover loss by district *d* and year *t*. *X_dt_* is a vector of biophysical controls and *W* indicates their spatial lags. *GP* is a vector of global commodity prices by year *t. NE* is a vector of conditions of Vietnam and neighboring and regional economies such as Laos, Cambodia, Myanmar, China, etc., including GDP per capita, GDP growth, and trade liberalization by trade as a percentage of GDP. *a* is the constant and *ε_dt_* is the error term. Variables were tested for normal distribution using a Shapiro-Wilks test and abnormally distributed variables were log-transformed, retested, and skew-corrected if necessary (see raw variable distributions in Figure 9 and 10 histograms for those that required skew correction). This model skeleton was thus used for linear, log-linear, and log-log specifications.

We used temporal lags of one to two years where relevant in order to account for the time delay between a change in price or economic conditions and the resulting effect on deforestation. We also further expound upon our hypotheses for relationships between these variables and deforestation in Table 1. A simple spatial lag of x model with OLS linear estimation was used due to the time series nature of the global and regional trend variables. Without a site-level fluctuation, methods such as fixed effects were not usable. As such, this estimation is subject to limitations discussed in the following section but nonetheless provides a useful baseline for understanding key commodities and regional economies’ effects on CAL deforestation.

Next, the more immediate social and economic determinants of deforestation were estimated using the following model. With district level data from 2000-2017, we used a site fixed-effect to account for time-invariant unobservables between districts. To select fixed effects over random effects we performed a Hausman test (threshold p<0.05) without clustered standard errors and a Sargan-Hansen test (threshold p<0.05) with clustered standard errors, both of which confirmed fixed over random effects. Because the panel is unbalanced due to missing data for certain districts in certain years, we were unable to perform space-time fixed effects and our sample size was reduced. However, like the underlying conditions model, we employed temporally lagged variables to account for time delay in effect on deforestation where relevant.

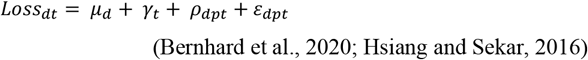

*μ_d_* is the sector fixed effect, *γ_t_* is time (year). *ρ_dpt_* is a vector of time-varying socioeconomic conditions at district level *d*, and provincial level *p*, including poverty rates, education levels, proxy for off-farm wages (proximity to urban area), proxy for human development (nighttime lights) at time *. Time invariant biophysical controls such as slope and elevation are not included in this term as the fixed effect accounts for those differences, but temperature and precipitation are included as they are time variant. *ε_dpt_* denotes the error term, unobservables across districts and provinces.

### 2.3 Scenarios of economic and socioeconomic conditions

We use the results from regression analysis in Phase 1 for scenario analysis. There are a number of different strategies that could be employed for scenario modeling, including forecasting. For the purposes of this baseline study, we began with simple methods for scenario analysis, using predictive margins based on fixed explanatory variable values. As is discussed further in the Results section, the following variables were identified as key determinants of deforestation: Chinese GDP and economic growth, rubber prices, poverty and/or local level of development at the district level, and education. Based on the Phase 1 models selected in the Results section, we fixed the values of these four variables to a set level using predictive margins, to determine how that variable’s effect on tree cover loss would change from the original model.

Our four key interest areas identified by Phase 1 modeling are displayed in Table 2. Table 2, which is also readapted in Table 13 to show how models were fit with fixed values for the variable of interest. Fixed values were selected using the existing historical minimum, maximum, and mean values for each raw variable. We followed the most successful model utilizing each variable, prioritizing R^2^ values. *X* is a vector of non-interest variables (the remainder of variables run normally in the regression) for a log-log spatial OLS. *ε_dpt_* is the error term for district *d*, province *p*, and at time *t*.

**Table 4.** Results of initial specification assessing the role only of biophysical control variables. Forest cover loss is the response variable. Spatial lags of slope and temperature are excluded due to collinearity with other biophysical lags. *** p<0.01, **p<0.05, *p<0.1. Robust standard errors in parentheses (clustered to district).

| **Tree cover loss** | **M1**<br>**Log-Linear** | **M2**<br>**Log-log** | **M2**<br>**Linear, unlagged** | **M3**<br>**Linear, lagged** |
| --- | --- | --- | --- | --- |
| *Elevation* | -0.0000664<br>(0.0003499) | 0.4358527<br>(0.3089644) | -61.48903<br>(994.8899) | 564.8219<br>(3202.348) |
| *Elevation (spatial lag)* | -0.0003219<br>(0.0003549) | -0.0007396<br>(0.0007395) |  | -675.0184<br>(2251.421) |
| *Slope* | -0.1505185***<br>(0.0193649) | -0.6848749**<br>(0.3196085) | -143393.2**<br>(68268.06) | 136226.5<br>(144568.4) |
| *Precipitation* | -0.0022168**<br>(0.0010802) | -0.364951<br>(0.2822432) | -12917.43***<br>(2729.952) | -15838.73**<br>(4485.264) |
| *Precipitation (spatial lag)* | 0.0018992<br>(0.0019875) | 0.0062531<br>(0.0037532) |  | 10683.24<br>(10280.5) |
| *Temperature* | 0.1395395***<br>(0.0308188) | 2.761337**<br>(1.36159) | 344775.5***<br>(79448.26) | 368600.3**<br>(150707.5) |
| *Existing forest cover* | 0.0001128***<br>(0.0000143) | 0.5519738***<br>(0.095402) | 305.9007***<br>(65.50001) | 305.54**<br>(137.344) |
| *Year* | 0.1540747***<br>(0.0114614) | 0.1599431<br>(0.01125) | 566752.9***<br>(44585) | 572376.1***<br>(62757.12) |
| *Proportion in protected area* | -0.0004213**<br>(0.0001436) | -0.0001883<br>(0.0002434) | -1106.464*<br>(627.2555) | -932.3636<br>(1065.383) |
| *Constant* | -299.4455***<br>(22.90498) | -320.0846***<br>(20.61693) | -1.14e+09***<br>(8.93e+07) | -1.16e+09***<br>(1.25e+08) |
| *n* | 680 | 653 | 680 | 680 |
| *R-sq* | 0.3809 | 0.5335 | 0.3033 | 0.3064 |

**Table 5.** Results of initial specification assessing the role types of agricultural croplands. Tree cover loss is the response variable. Spatial lags of slope and temperature are excluded due to collinearity with other biophysical lags. *** p<0.01, **p<0.05, *p<0.1. Robust standard errors in parentheses (clustered to district).

| **Tree cover loss** | **M1**<br>**Linear** | **M2**<br>**Linear** | **M3**<br>**Log-log** | **M4**<br>**Log-log** |
| --- | --- | --- | --- | --- |
| *Irrigated cropland* | -833.3261<br>(906.9169) | 2077.859<br>(1899.037) | 0.201797***<br>(0.0556102) | -0.0183835<br>(0.0588528) |
| *Irrigated cropland*<br>*(spatial lag)* | -9644.399***<br>(1377.468) | 2453.251<br>(2151.895) | -0.000017 (0.0003011) | -0.0030648***<br>(0.0004495) |
| *Mosaic cropland* | 877.1***<br>(230.8801) | 1064.529**<br>(371.0898) | 0.6772165***<br>(0.0374728) | 0.4813517***<br>(0.0523451) |
| *Rainfed cropland* | 945.9721**<br>(319.3324) | 201.0895<br>(772.5546) | 0.3115648***<br>(0.0708857) | 0.0315687<br>(0.0737679) |
| *Year* | 563692.1***<br>(42369.38) | 558299***<br>(55657.13) | 0.1547289***<br>(0.0084733) | 0.1612482***<br>(0.0095476) |
| *Elevation* | 2023.826<br>(1733.584) |  |  | 0.0005811**<br>(0.000315) |
| *Elevation (spatial lag)* | -6371.642***<br>(1270.518) |  |  | -0.0025902***<br>(0.0003977) |
| *Slope* | 303864.9***<br>(65659.05) |  |  | 0.1181852***<br>(0.018947) |
| *Precipitation* | -15802.01***<br>(3180.814) |  |  | -0.0033265***<br>(0.0008038) |
| *Precipitation (spatial lag)* | 6710.539<br>(7024.947) |  |  | -0.0010668<br>(0.0018185) |
| *Temperature* | 582777.2***<br>(106160.1) |  |  | 0.1334604***<br>(0.0373387) |
| *Existing forest cover* | 187.5387**<br>(64.55037) | 297.2574**<br>(95.79903) | 0.0000329**<br>(0.0000119) | 0.0000303**<br>(0.0000146) |
| *Proportion in protected area* | -2600.75***<br>(737.3091) | 1795.763<br>(1412.523) | -0.0000232 (0.000146) | -0.0005778***<br>(0.0001547) |
| *Constant* | -1.14e+09***<br>(8.49e+07) | 1.12e+09***<br>(1.11e+08) | -299.7585***<br>(16.98636) | -314.0616***<br>(18.92626) |
| *n* | 680 | 757 | 718 | 656 |
| *R-sq* | 0.3921 | 0.3437 | 0.5531 | 0.6000 |

**Table 6.** Results of the Underlying Causes model, for macroeconomic conditions that may drive deforestation. Forest cover loss is the response variable. Square brackets indicate temporal lag (e.g., [L1] = one year time lag). Parentheses in variable name indicate natural log or skew correction. M1 and M2 are economies specifications (trade was excluded from M2 as all were collinear with the inclusion of temporal lags), while M3 to M5 are price models. Variables omitted due to collinearity are indicated with “omit,” while blank cells indicate that that variable was not included in the specification due to previously identified collinearity or for the purposes of a given specification. The same values of biophysical variables and control variables between M1 and M2 and between M3 and M5 should be explored in further research. *** p<0.01, **p<0.05, *p<0.1. Robust standard errors in parentheses (clustered to district). Square-meter values are converted into hectares in the interpretation.

| Tree cover loss | M1<br>NRE with trade | M2<br>NRE temporal<br>lags, no trade | M3<br>Prices | M4<br>Prices and<br>temporal lags | M5<br>Prices and<br>temporal lags |
| --- | --- | --- | --- | --- | --- |
| *VN per capita GDP* | 19994.16**<br>(11262.26) | omit | -816.4136<br>(874.4272) |  |  |
| *VN per capita GDP [L1]* |  | 4951.827<br>(13162.37) |  | 5610.994***<br>(707.564) |  |
| *VN trade as component GDP* | 208987.8***<br>(45761.97) |  |  |  |  |
| *China per capita GDP* | 4264.225<br>(7058.629) | 39629.34***<br>(5493.504) |  |  |  |
| *China per capita GDP [L1]* |  | 17249.27**<br>(5433.028) |  |  |  |
| *China trade as component GDP* | 207368.9<br>(180703.3) |  |  |  |  |
| *Lao per capita GDP* | -55682.33**<br>(20717.61) | -11798.15<br>(9308.188) |  |  |  |
| *Lao per capita GDP [L1]* |  | 35640.25***<br>(8916.897) |  |  |  |
| *Lao trade as component GDP* | 318663.6**<br>(96782.06) |  |  |  |  |
| *Cambodia per capita GDP* | 20311.78**<br>(8304.602) | omit |  |  |  |
| *Cambodia per capita GDP [L1]* |  | 12010.87<br>(6702.015) |  |  |  |
| *Cambodia trade as component GDP* | omit |  |  |  |  |
| *Thai per capita GDP* | 2739.385*<br>(1538.123) | 7083.602<br>(2386.173) |  |  |  |
| *Thai per capita GDP [L1]* |  | 2219.646<br>(1281.933) |  |  |  |
| *Thai trade as component GDP* | 51391.73**<br>(20270.9) |  |  |  |  |
| *Thai trade as component GDP [L1]* | omit |  |  |  |  |
| *Myanmar per capita GDP* | 9773.863<br>(10102.32) | -45919.89***<br>(8489.44) |  |  |  |
| *Myanmar per capita GDP [L1]* |  | omit |  |  |  |
| *Myanmar trade as component GDP* | 637335.5***<br>(144350.7) |  |  |  |  |
| *India per capita GDP* | 10479.59**<br>(3270.766) | 61132.85***<br>(8880.916) |  |  |  |
| *India per capita GDP [L1]* |  | 16821.04**<br>(8180.191) |  |  |  |
| *India trade as component GDP* | omit |  |  |  |  |
| *Japan per capita GDP* | -7899.611***<br>(1796.776) | -2277.945**<br>(1017.623) |  |  |  |
| *Japan per capita GDP [L1]* |  | -889.8979<br>(572.7266) |  |  |  |
| *Japan trade as component GDP* | omit |  |  |  |  |
| *Rice price (ln k)* |  |  | -43332.92***<br>(7706.772) |  | -24900.05***<br>(3385.241) |
| *Rice price (ln k) [L1]* |  |  |  | 10280.74**<br>(5020.901) |  |
| *Hardwood price index (ln)* |  |  |  |  | 497920.3***<br>(87459.06) |
| *Hardwood price index (ln) [L1]* |  |  | omit |  |  |
| *Softwood price index* |  |  |  | omit |  |
| *Softwood price index [L1]* |  |  |  |  | omit |
| *Metals price index (ln k)* |  |  |  |  | 164161.6*<br>(83019.28) |
| *Metals price index (ln k) [L1]* |  |  | omit |  | omit |
| *Rubber price* |  |  | 27962.46***<br>(6975.312) |  | 104805.3**<br>(48247.48) |
| *Rubber price [L1]* |  |  |  | 78992.96***<br>(11961.03) |  |
| *Tea coffee cocoa price index* |  |  | 115584.3***<br>(21341.72) |  | -18657.42<br>(15094.74) |
| *Tea coffee cocoa price index [L1]* |  |  |  |  | omit |
| *Cereals price index* |  |  | 131355.4***<br>(27119.04) |  |  |
| *Cereals price index [L1]* |  |  |  | -15861.78<br>(14451.38) |  |
| *Poverty rate* |  |  | 154409.4<br>(93337.66) | 231338**<br>(74927.3) | 154409.4<br>(93337.66) |
| *Irrigated cropland* |  |  | 4024.161*<br>(2348.825) | 2341.974<br>(2139.536) | 4024.161*<br>(2348.825) |
| *Urban area* | -8833.604<br>(16563.66) | -8833.604<br>(16563.66) | -18960.44<br>(15022.94) | -9331.049<br>(15196.83) | -18960.44<br>(15022.94) |
| *District contribution to GDP* | -1378815**<br>(560398.4) | -1378815**<br>(560398.4) |  |  |  |
| *Elevation* | 1985.772<br>(3046.041) | 1985.772<br>(3046.041) |  | -1909.749<br>(2107.021) |  |
| *Elevation (spatial lag)* | -1722.745<br>(2218.22) | -1722.745<br>(2218.22) | -1985.333<br>(2140.133) |  | -1985.333<br>(2140.133) |
| *Slope* | -57185.41<br>(145566.2) | -57185.41<br>(145566.2) | -1985.333<br>(2140.133) | 588075.5**<br>(166730.3) | -1985.333<br>(2140.133) |
| *Precipitation* | 1060.3<br>(5574.843) | 1060.3<br>(5574.843) |  | 19347.79<br>(15543.34) |  |
| *Precipitation (spatial lag)* | -2063.274<br>(10643.47) | -2063.274<br>(10643.47) | 21717<br>(16820.62) |  | 21717<br>(16820.62) |
| *Temperature* | 391184.4**<br>(144237.4) | 391184.4**<br>(144237.4) | 200748.7<br>(241258.6) |  | 200748.7<br>(241258.6) |
| *Existing forest coverage* | 229.3596*<br>(132.8239) | 229.3596*<br>(132.8239) |  |  |  |
| *Proportion in protected area* | -624.7059<br>(1292.044) | -624.7059<br>(1292.044) | 960.1925<br>(1589.235) | 1089.264<br>(1491.94) | 960.1925<br>(1589.235) |
| *Constant* | 1.32e+08**<br>(5.39e+07) | -8.99e+07***<br>(1.77e+07) | 2.01e+07**<br>(7348026) | 2.30e+07***<br>(6010935) | 2.01e+07**<br>(7348026) |
| *n* | 680 | 680 | 0.3433 | 0.2984 | 0.3433 |
| *R-sq* | 0.3822 | 0.3823 | 305 | 256 | 305 |

**Table 7.**
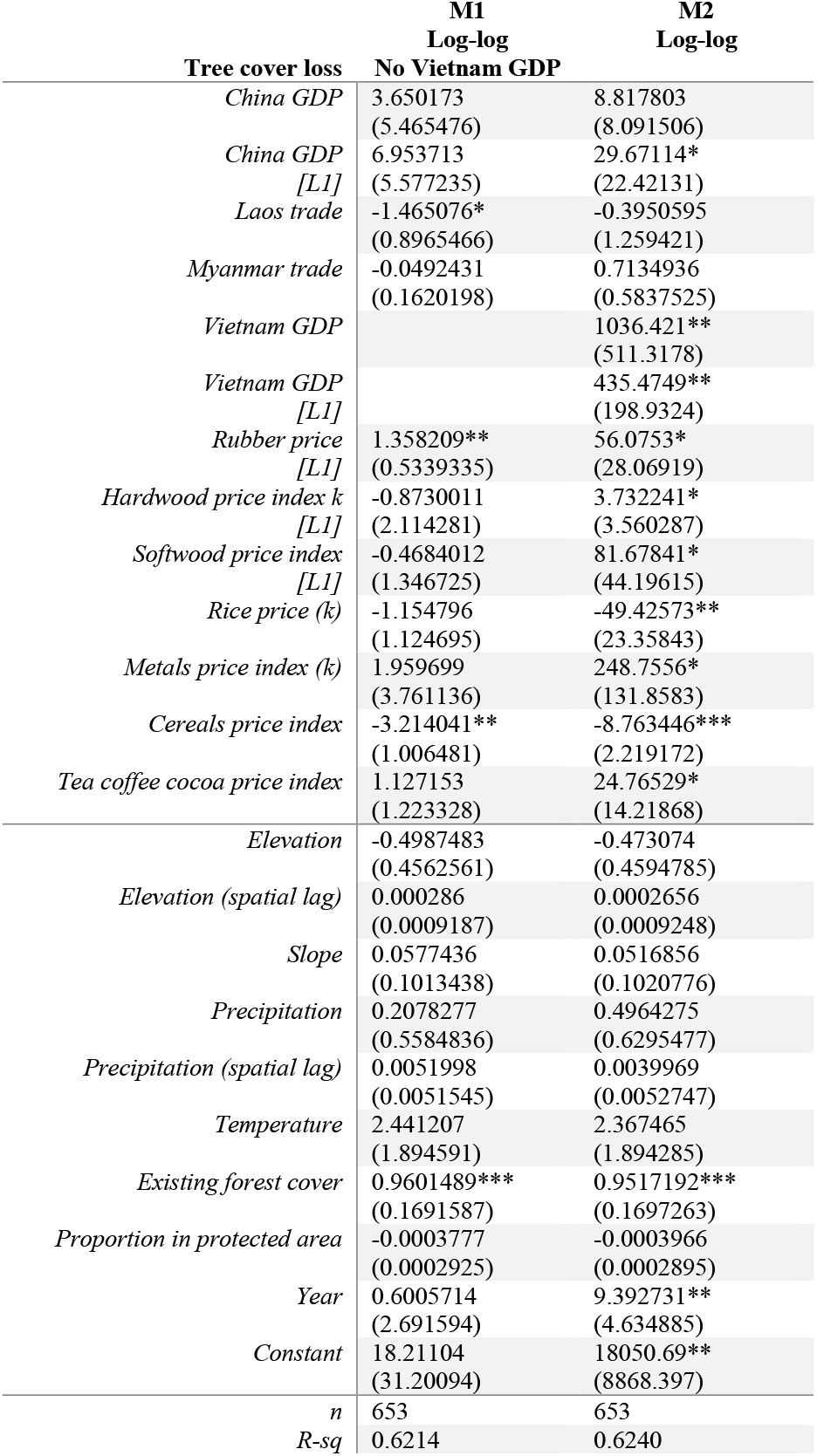
Restricted model for Underlying Causes, for macroeconomic conditions that may drive deforestation. This restricted specification includes key variables identified from Table 6 and is based on the best specification from Tables 4 and 5. This table is thus log-log only, based on the R^2^ values from Table 4 M2 and Table 5 M4. Forest cover loss is the response variable. Square brackets indicate temporal lag (e.g., [L1] = one-year time lag). Parentheses in variable name indicate natural log or skew correction. M1 and M2 are economies models, while M3 to M5 are price models. *** p<0.01, **p<0.05, *p<0.1. Robust standard errors in parentheses (clustered to district).

**Table 8.** Results of the Immediate Causes model, for micro and socioeconomic conditions that may drive deforestation. Forest cover loss is the response variable. Fixed effects were selected over random effects following a Hausmann test (p=0.0087) without clustered standard errors and a Sargan-Hansen test with clustered standard errors. Variables omitted due to collinearity are indicated with “omit,” while blank cells indicate that that variable was not included in the specification due to previously identified collinearity or for the purposes of a given specification. *** p<0.01, **p<0.05, *p<0.1. Robust standard errors in parentheses (clustered to district).

| **Tree cover loss** | **M1** | **M2**<br>**Temporal lags** | **M3**<br>**(No FE)** | **M4**<br>**(No FE)** |
| --- | --- | --- | --- | --- |
| *Population density (spatial lag)* | -35341.38<br>(33808) | -29057.39<br>(34522.63) | -363.88<br>(673.7428) | -801.613<br>(1047.978) |
| *Poverty rate (categorical)* | 365447.3*<br>(45168.57) |  |  |  |
| *District contribution to GDP [L1]* | Omit |  | 172395.4***<br>(529051) | 1044925***<br>(233935.9) |
| *Secondary school graduation ratio* | -126960.5*<br>(74337.03) |  |  |  |
| *Secondary school graduation ratio [L1]* |  | -36425.6<br>(31534.99) | -29339.55***<br>(21921.96) | -124201.5***<br>(23544.69) |
| *Irrigated cropland* | 86690<br>(126354.4) |  |  |  |
| *Irrigated cropland [L1]* |  | 221580.3<br>(152534.1) | 475.4306<br>(921.3167) | 1860.187<br>(1317.673) |
| *Mosaic cropland* | 22678.12**<br>(7209.053) |  |  |  |
| *Mosaic cropland [L1]* |  | 29060.38**<br>(9003.066) | 2036.557**<br>(579.7422) | 2142.061**<br>(609.2791) |
| *Rainfed cropland* | 16745.82<br>(44247.01) |  |  |  |
| *Rainfed cropland [L1]* |  | 60999.82*<br>(34972.42) | 370.7381***<br>(326.2099) | 237.5954***<br>(300.1541) |
| *Nighttime lights (spatial lag) [L1]* | Omit | Omit | -5133.074<br>(49774.15) | 260478.3<br>(194932.1) |
| *Precipitation (spatial lag)* | Omit | Omit | 5620.537*<br>(4219.364) | 3653.445<br>(8258.201) |
| *Temperature (spatial lag)* | 1664024*<br>(758860) | 494880<br>(577849.2) | 56545.33*<br>(139695.1) | 79941.23<br>(196275.9) |
| *Constant* | -7.52e+07<br>(5.29e+07) | -1.20e+07<br>(4.60e+07) | 3024997<br>(4058081) | -3856935<br>(4769536) |
| *n* | 143 | 143 | 143 | 437 |
| *Within R-sq* | 0.2994 | 0.2885 |  |  |
| *R-sq* |  |  | 0.2892 | 0.3413 |

**Table 9.**
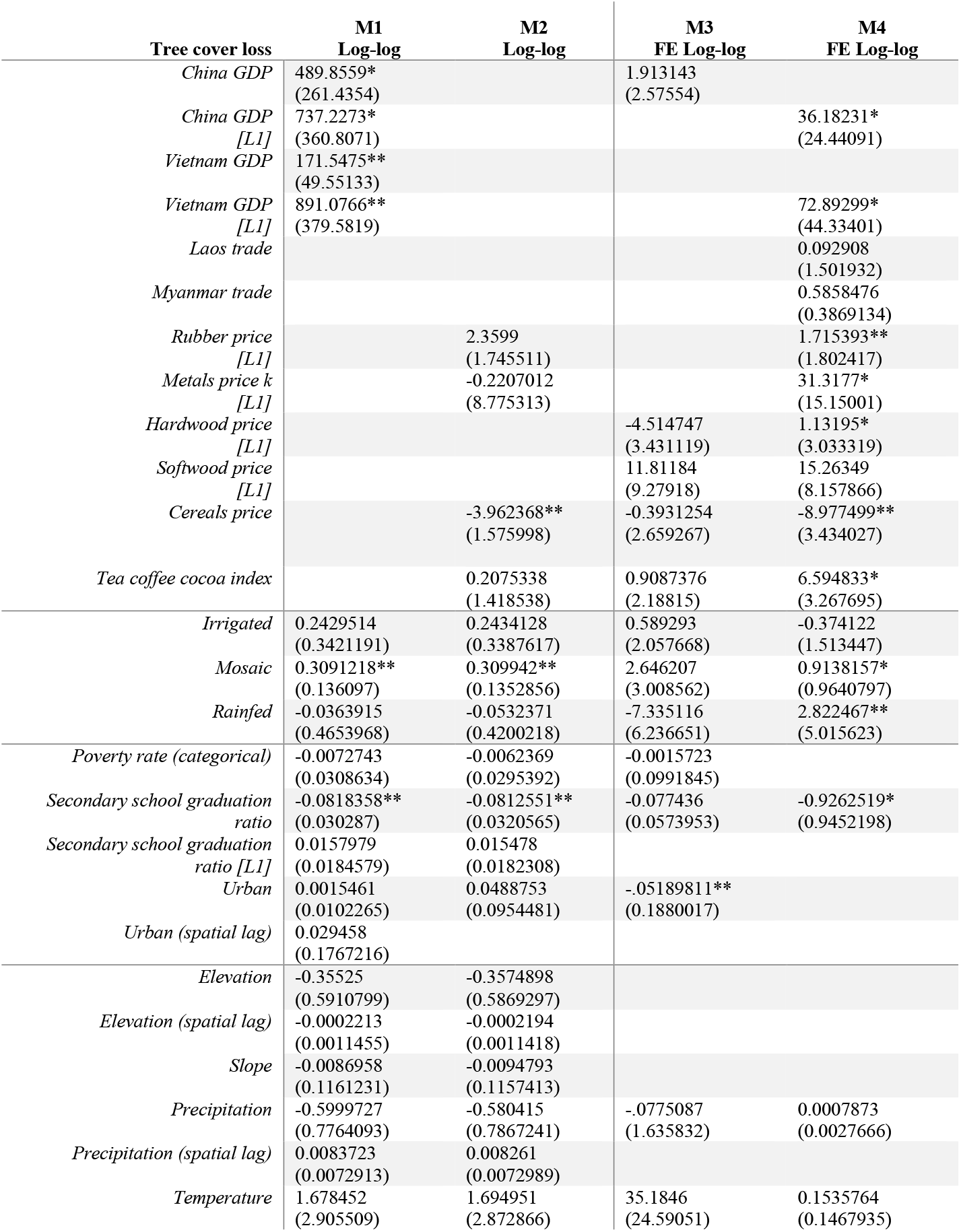
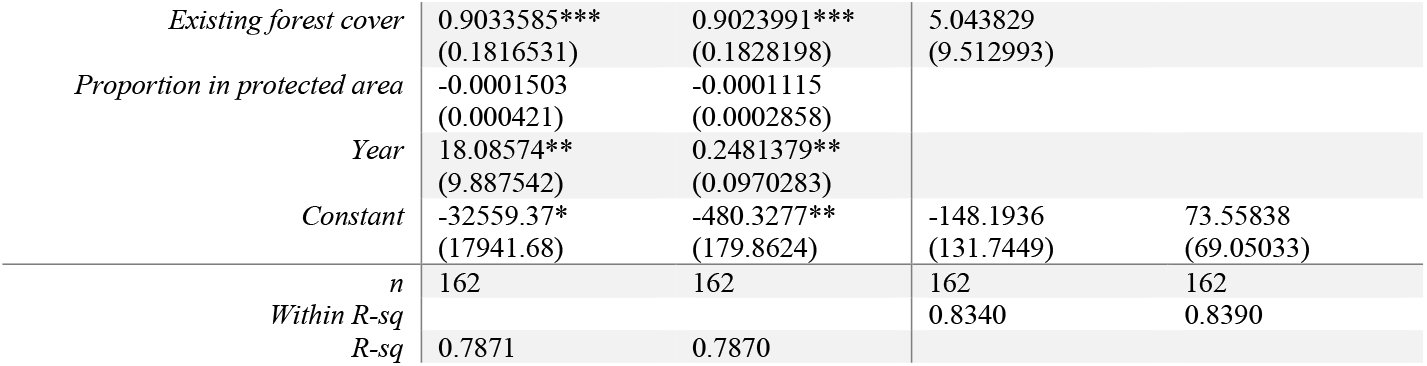
Results of a combined restricted model, using key variables identified from the results of both underlying and immediate determinants models. Economies variables and price variables are not included in the same specifications. Forest cover loss is the response variable. For M3/M4, fixed effects were selected over random effects following a Hausmann test (p=0.0087) without clustered standard errors and a Sargan-Hansen test with clustered standard errors. Time-invariant variables (such as elevation and slope) are excluded. Low statistical significance is expected in M3/M4 due to low sample size for FE. *** p<0.01, **p<0.05, *p<0.1. Robust standard errors in parentheses (clustered to district).

**Table 10.**
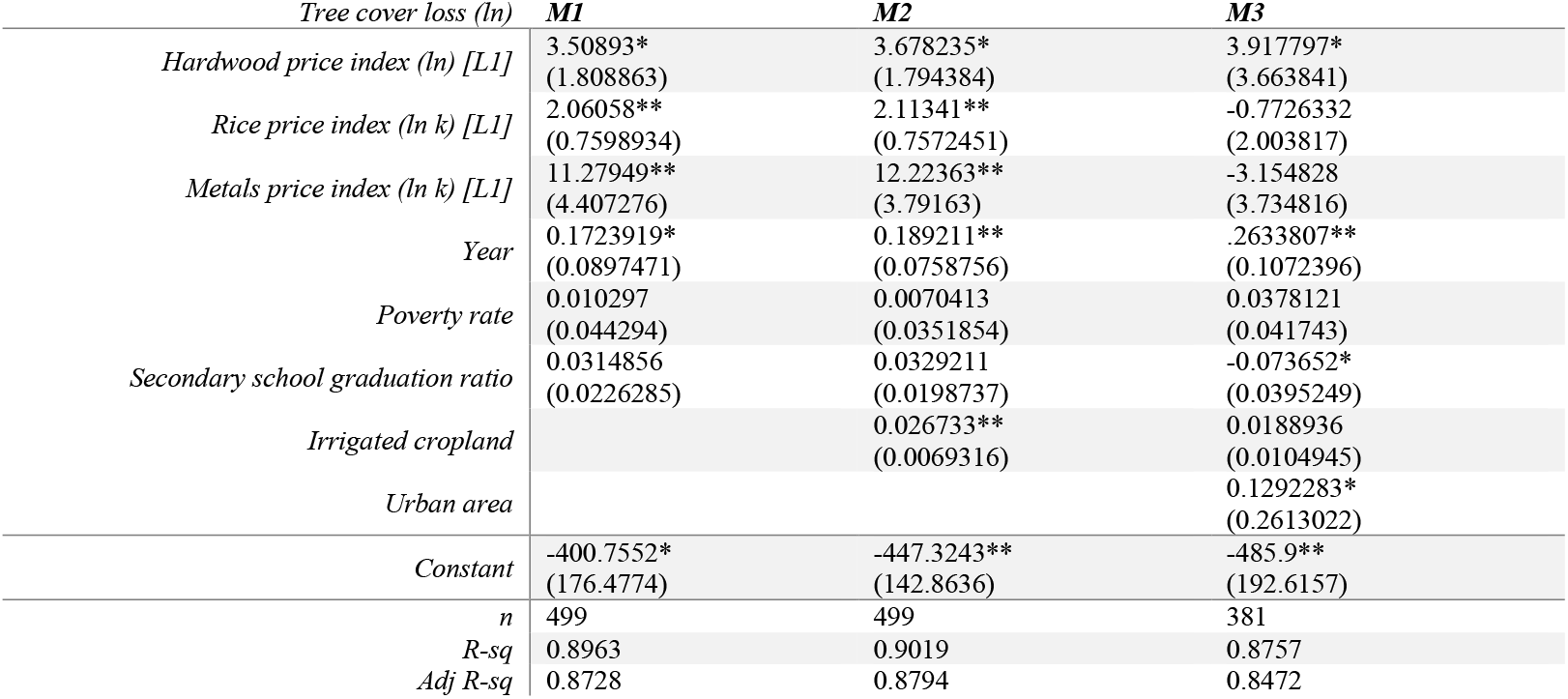
Results for spatially unlagged (spatial) regression specifications combining underlying and immediate determinants. This was an exploratory specification to observe variables prior to adjustment for spatial autocorrelation and to observe combined effect of global prices and district level immediate determinants of deforestation. These results are strictly correlative as coefficients are biased by spatial autocorrelation and important biophysical variables are not included. *** p<0.01, **p<0.05, *p<0.1. Robust standard errors in parentheses (clustered to district).

**Table 11.**
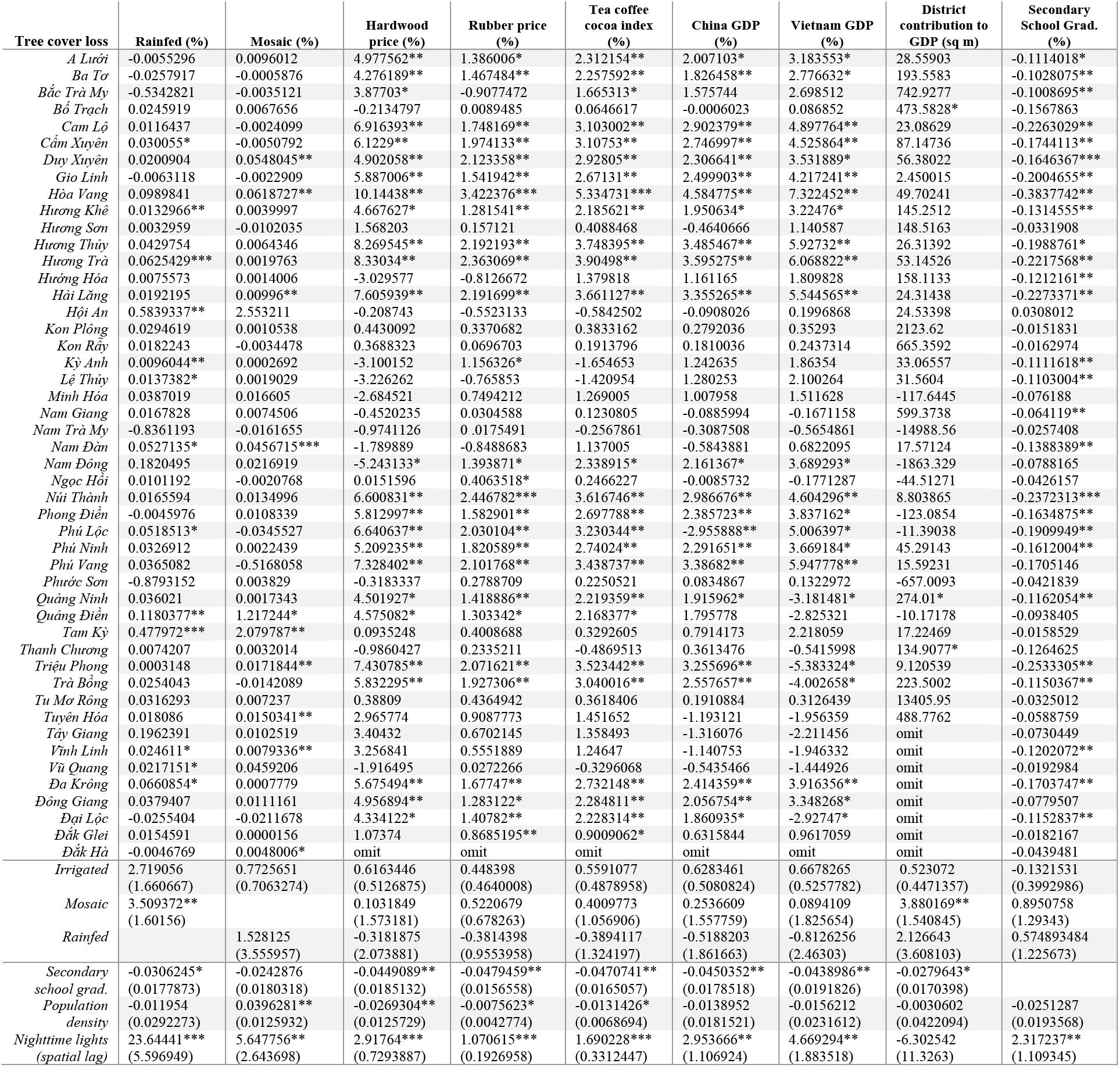
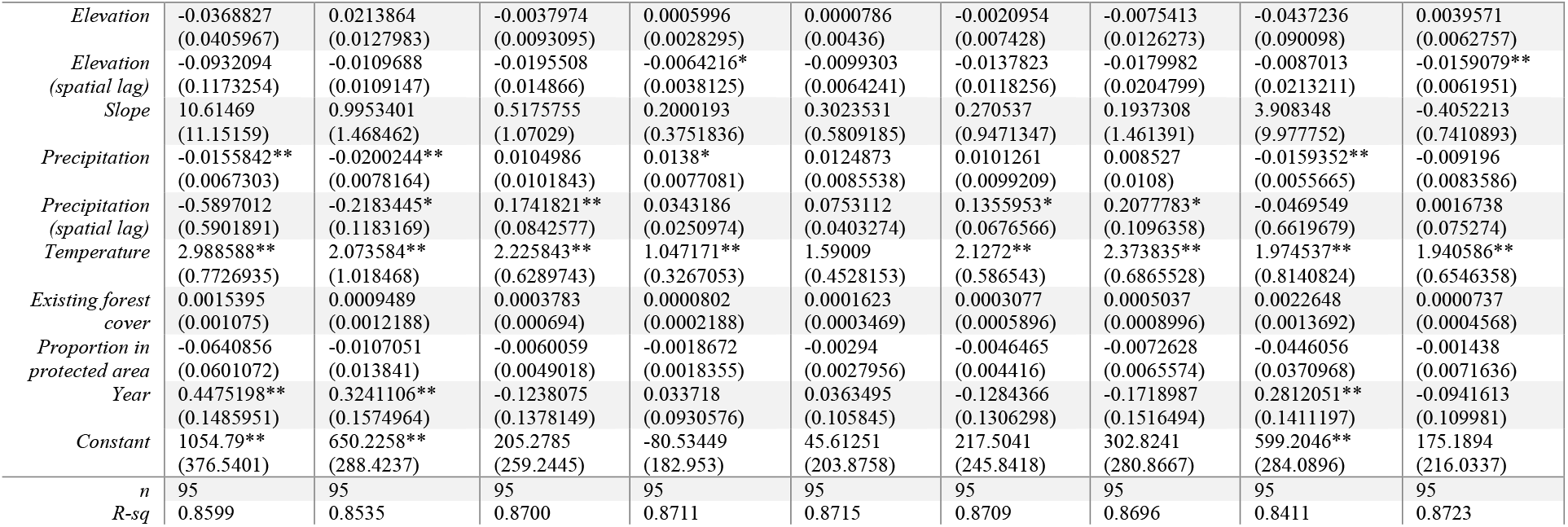
Based on the model and results of Table 9, we ran a district-specific model, using a Tier 2/district factor variable (interaction) for each key explanatory variable identified in Table 9. The corresponding maps for each district’s deforestation “sensitivity” to the economic variable (the magnitude of the interaction coefficient) are shown in Figures 20 and 21. Districts with insufficient data for estimation are not included. *** p<0.01, **p<0.05, *p<0.1. Robust standard errors in parentheses (clustered to district).

**Table 12.**
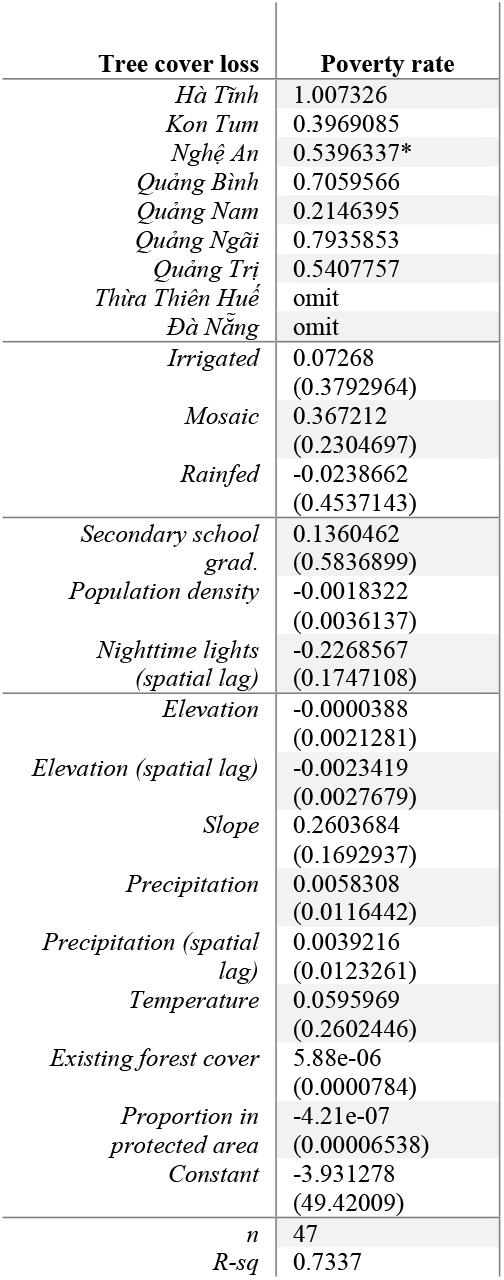
Based on the model and results of Table 9, we ran a province-specific model, using a Tier 1/province factor variable (interaction) for poverty rate. This done at the Tier 1 level, rather than Tier 2 (as in Table 11) because the poverty rate data was available only at the province level. The corresponding map for each province’s deforestation “sensitivity” to poverty rate (the magnitude of the interaction coefficient) is shown in Figure 20, bottom left. *** p<0.01, **p<0.05, *p<0.1. Robust standard errors in parentheses (clustered to province).

| **Tree cover loss** | **Poverty rate** |
| --- | --- |
| *Hà Tĩnh* | 1.007326 |
| *Kon Tum* | 0.3969085 |
| *Nghệ An* | 0.5396337* |
| *Quảng Bình* | 0.7059566 |
| *Quảng Nam* | 0.2146395 |
| *Quảng Ngãi* | 0.7935853 |
| *Quảng Trị* | 0.5407757 |
| *Thừa Thiên Huế* | omit |
| *Đà Nẵng* | omit |
| *Irrigated* | 0.07268<br>(0.3792964) |
| *Mosaic* | 0.367212<br>(0.2304697) |
| *Rainfed* | -0.0238662<br>(0.4537143) |
| *Secondary school grad.* | 0.1360462<br>(0.5836899) |
| *Population density* | -0.0018322<br>(0.0036137) |
| *Nighttime lights (spatial lag)* | -0.2268567<br>(0.1747108) |
| *Elevation* | -0.0000388<br>(0.0021281) |
| *Elevation (spatial lag)* | -0.0023419<br>(0.0027679) |
| *Slope* | 0.2603684<br>(0.1692937) |
| *Precipitation* | 0.0058308<br>(0.0116442) |
| *Precipitation (spatial lag)* | 0.0039216<br>(0.0123261) |
| *Temperature* | 0.0595969<br>(0.2602446) |
| *Existing forest cover* | 5.88e-06<br>(0.0000784) |
| *Proportion in protected area* | -4.21e-07<br>(0.00006538) |
| *Constant* | -3.931278<br>(49.42009) |
| *n* | 47 |
| *R-sq* | 0.7337 |

**Table 13.** Results for estimating forest cover loss using fixed values (predictive margins) for Chinese economic growth, rubber price fluctuation, poverty reduction, and secondary school graduation results for each possible outcome is indicated with green, red, or yellow.

|  |  |
| --- | --- |
| **Chinese economic growth**<br>$\ln (Loss)_{dt} = \alpha + \beta_1 \ln [X]_{dt} + \beta_2 WX_{dt} + \beta_x \ln [China]_t + \varepsilon_{dt}$<br>Table 7 M1<br><u>Possible outcomes:</u><br>1. **Economic contraction** (*China* = 7.983889)<br>2. **Maintained growth** (*China* = 8.912644)<br>3. **Economic boom** (*China* = 9.729535) | **Rubber price fluctuation**<br>$\ln (Loss)_{dt} = \alpha + \beta_1 \ln [X]_{dt} + \beta_2 WX_{dt} + \beta_x \ln [Rubber]_t + \varepsilon_{dt}$<br>Table 7 M2<br><u>Possible outcomes:</u><br>1. **Drop in price** (*Rubber* = 3.261404)<br>2. **Increase in price** (*Rubber* = 5.386853) |
| **Poverty reduction**<br>$Loss_{dpt} = \alpha + \beta_1 X_{dpt} + \beta_2 WX_{dpt} + \beta_x Pov_t + \varepsilon_{dpt}$<br>Table 6 M4<br><u>Possible outcomes:</u><br>1. **Worsening poverty** (*Pov* = 2)<br>2. **Maintain current levels** (*Pov* = 3)<br>3. **Continued improvement** (*Pov* = 5)<br>(poverty rate quintiles) | **Education level improvement**<br>$Loss_{dpt} = \alpha + \beta_1 X_{dpt} + \beta_2 WX_{dpt} + \beta_x Edu_t + \varepsilon_{dpt}$<br>Table 8 M1<br><u>Possible outcomes:</u><br>1. **Current secondary graduation rates are maintained** (*Edu* = 91.42638)<br>2. **Graduation rates increase** (*Edu* = 99.86) |

Predictive margins are essentially a simple tool for answering “what if”-type questions without complex forecasting requiring high R^2^ values. Essentially, these margins enable controlling for compositional differences in the explanatory variables to be able to say, roughly, what the tree cover loss outcome could be in that case. For example, how would tree cover loss change in this model if rubber prices increased to their observed maximum? Or, how would tree cover loss change if all district secondary school graduation rates increased to the currently observed maximum? The predictive margins results are not interpretable alone and are be compared to the tree cover loss outcome in the original model on which the predictive margins are based.

### 2.4 Limitations

The methods used in this study are subject to standard issues in econometrics in addition to limitations specific to the uses of spatial data. Our dataset utilizes variables from a variety of sources, each with their own potential biases. Importantly, we note that tree cover loss data from (Hansen 2013) can be the result of a number of activities and is not always deforestation. However, we assume that tree cover loss is a reasonable proxy for human-driven deforestation. Next, while we have sought to use robust econometric techniques that account for omitted variables and endogeneity, we note that these specifications are imperfect and the underlying conditions model in particular is vulnerable to these issues due to its use of coarse time series data without spatial variation. Lastly, standard errors are unlikely to be entirely independent across observations as policy and local governance variation, in addition to many other factors, effects spatial distribution of deforestation within provinces and districts. Throughout the analysis, we explore clustering standard errors to province level (Tier 1) and district level (Tier 2) to ensure that they are robust to heteroscedasticity and take the more conservative estimate when relevant in terms of p-value. (Stock and Watson, 2006). Our Phase 2 scenario modeling is particularly sensitive to these issues, and the linearity of models run in this study, so these are to be interpreted with extra caution and as a baseline.

This dataset also relies on data from multiple scales, from global macroeconomic conditions and national level statistics to finer-scaled raster data for tree cover loss. Problems with this movement between scales, known as the “ecological fallacy,” are discussed by Moulton (1990). Conclusions drawn about individual units based on aggregated groups are problematic as phenomena occurring at a local scale cannot be fully understood by coarse-scale trends. However, our research design and framework intentionally move from coarse to fine in an effort to understand the different dimensions at each level, from underlying to immediate determinants of deforestation.

As will be discussed further throughout this study, we used available secondary data for this study, which established a baseline for future research. Future studies should utilize full GSOV household survey data for socioeconomic conditions at the district level or finer, for which this study was limited by available online downloads (see Appendix II, Data Availability Assessment for further discussion of this). Additionally, our results isolate limited causality but could be confirmed by identifying instrumental variables or “shock” events to enable use of quasi-experimental and natural experimental methods. Using Lao data could also enable a regression discontinuity design study to a similar effect.

## 3. Results

### 3.1 Identifying potential determinants of deforestation

#### 3.1.1 Exploratory spatial data analysis (ESDA)

ESDA revealed spatial autocorrelation of most key variables. While it is clear that biophysical variables would be spatially autocorrelated, as high elevation and high slope areas would be associated in mountainous regions of CAL, for example, identifying clustering of socioeconomic variables relative to deforestation is of standalone interest. Further, this spatial autocorrelation indicates that models failing to account for spatial sorting would be biased.

In addition to spatial autocorrelation, the correlation matrix in Table 3 and scatterplot matrices in Figures 9 and 10 further illustrate the collinearity of variables and bivariate relationships between key variables. Figures 11–16 show this clustering through coverage and statistical maps for several key variables, both biophysical and socioeconomic. Figures 17–19 present results of bivariate local Moran’s I for four potential immediate-scale socioeconomic determinants of deforestation, illustrating spatial association between these variables and deforestation in CAL equivalent to a simple spatial autoregression (see Appendix III). Our key finding from the bivariate statistics is that the hotspot areas of deforestation are not spatially collocated with population-dense, heavily economic areas. Evidence suggests that higher poverty rates and smaller scale subsistence agriculture are more likely to be driving deforestation at the immediate scale. Figure 17 demonstrates that while high poverty rates are spatially associated with tree cover loss (0.1684), population density does not display this relationship (−0.2661). This provides some evidence that population pressures in the forest frontier areas may not be a key spatial driver of deforestation, but rather immediate determinants of deforestation are more likely to be poverty and expansion of subsistence agriculture. To further support this finding, Figure 19 shows that neither nighttime lights nor irrigated cropland is spatially associated with tree cover loss (−0.3452, −0.1392, respectively). These findings are also consistent with the existing literature focusing on determinants of deforestation in Vietnam (Müller and Munroe, 2005).

**Figure 11.**
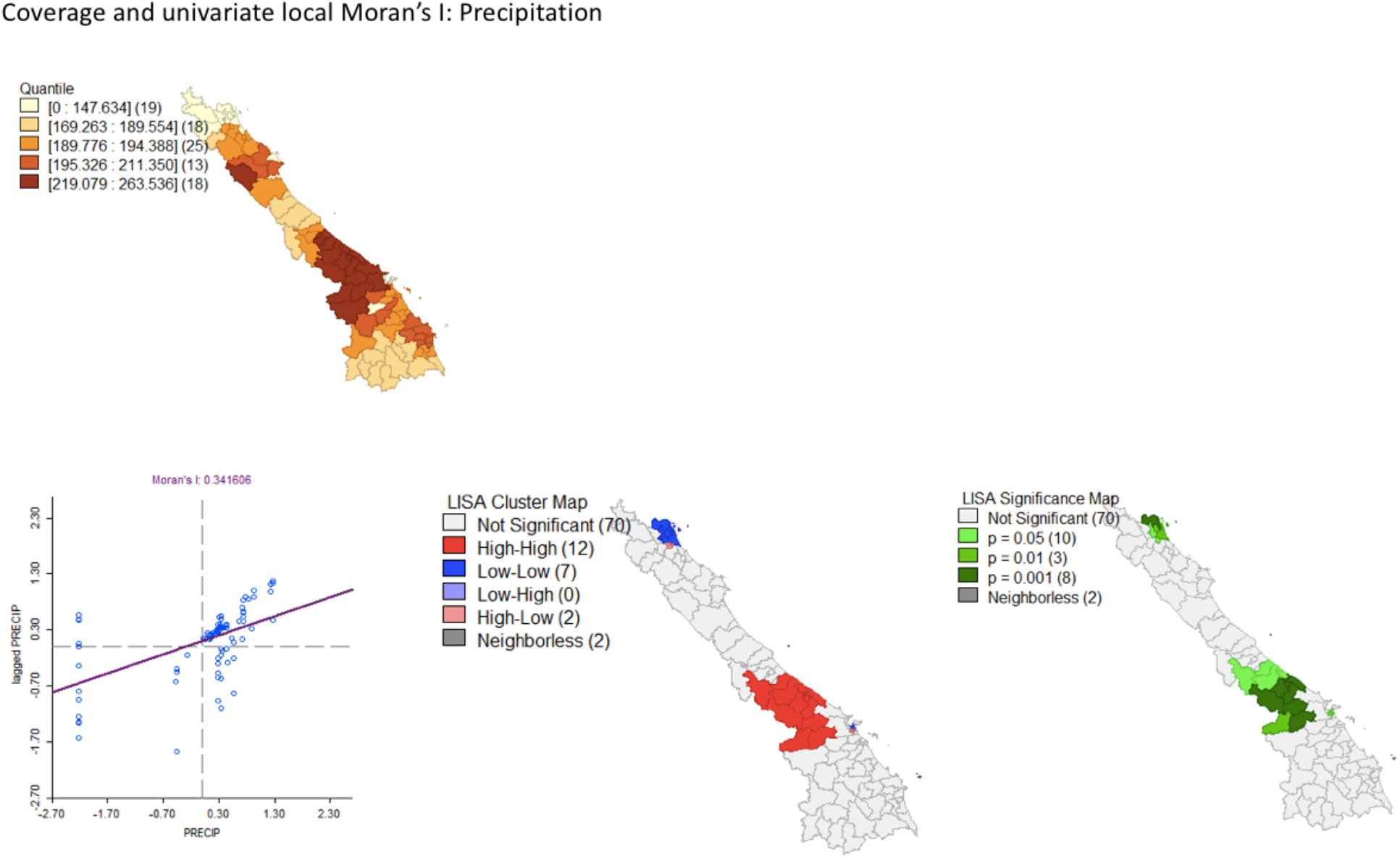
Mean coverage map by district and univariate local Moran’s I of precipitation in CAL. Displays significant spatial autocorrelation (Moran’s I: 0.3416) and so a spatially lagged precipitation variable was constructed.

**Figure 12.**
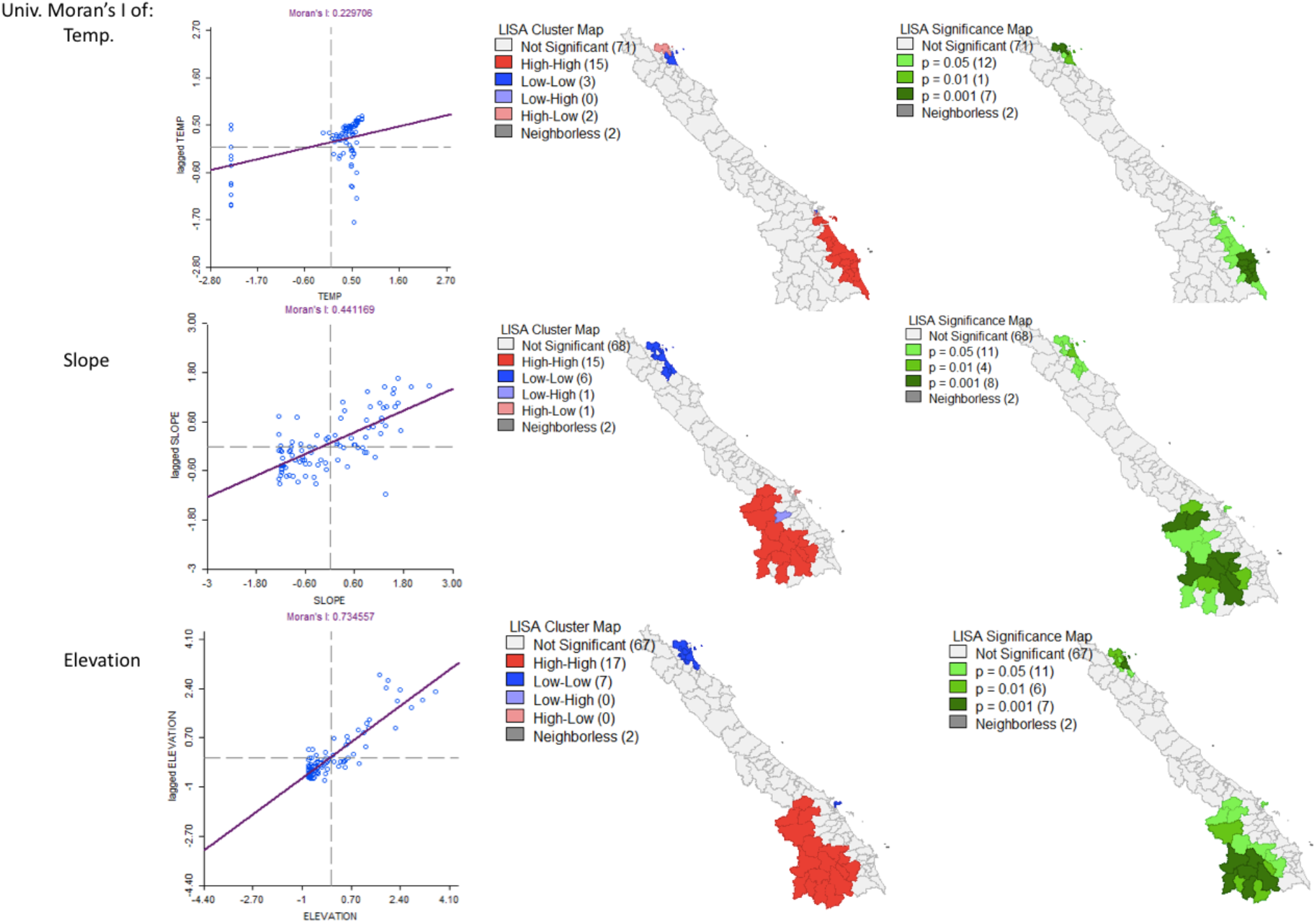
Univariate local Moran’s I of temperature, slope, and elevation in CAL. All display significant spatial autocorrelation (Moran’s I: 0.2297, 0.4411, 0.7346, respectively) and so a spatially lagged variables were constructed.

**Figure 13.**
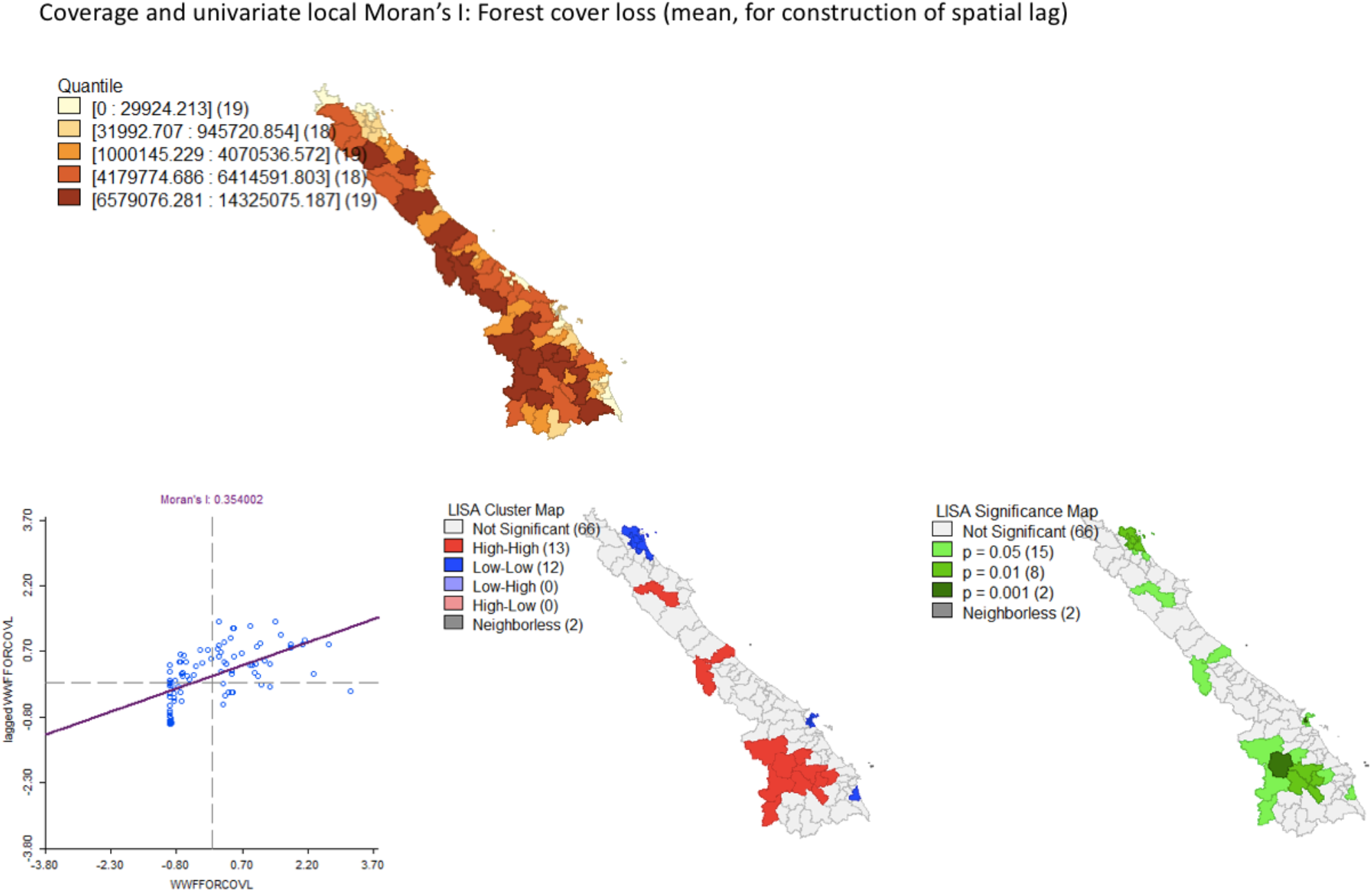
Mean coverage map by district and univariate local Moran’s I of forest cover loss in CAL. Averaged annually to construct the spatial weight. Displays significant spatial autocorrelation (Moran’s I: 0.354) and so a spatially lagged precipitation variable was constructed for spatially autoregressive modeling.

**Figure 14.**
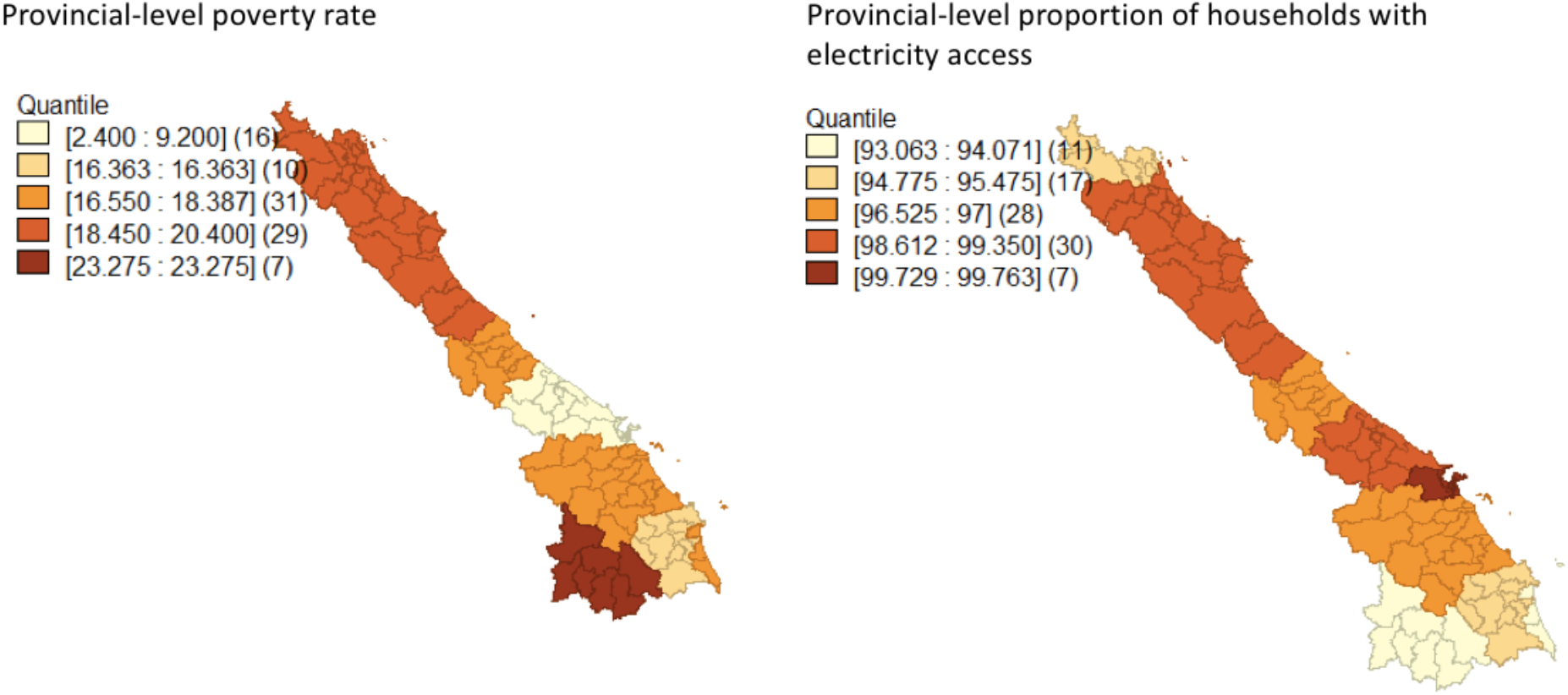
Mean coverage maps at the provincial level for poverty rates and proportion of households with electricity access. Used as categorical variable rather than continuous in modeling due to provincial level grouping by GSOV rather than district level.

**Figure 15.**
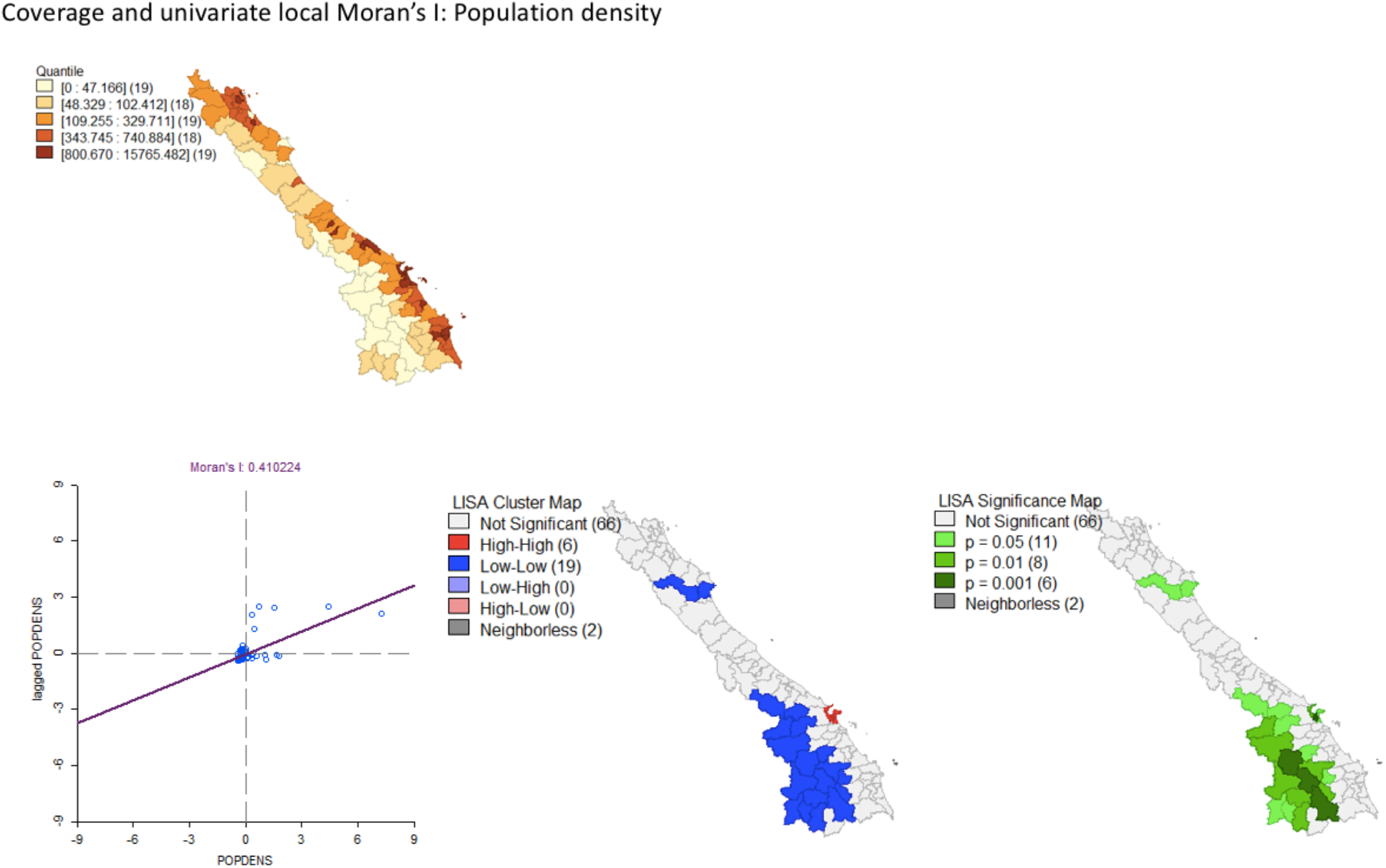
Mean coverage map by district and univariate local Moran’s I of population density in CAL. Displays significant spatial autocorrelation (Moran’s I: 0.4102) and so a spatially lagged population density variable was constructed.

**Figure 16.**
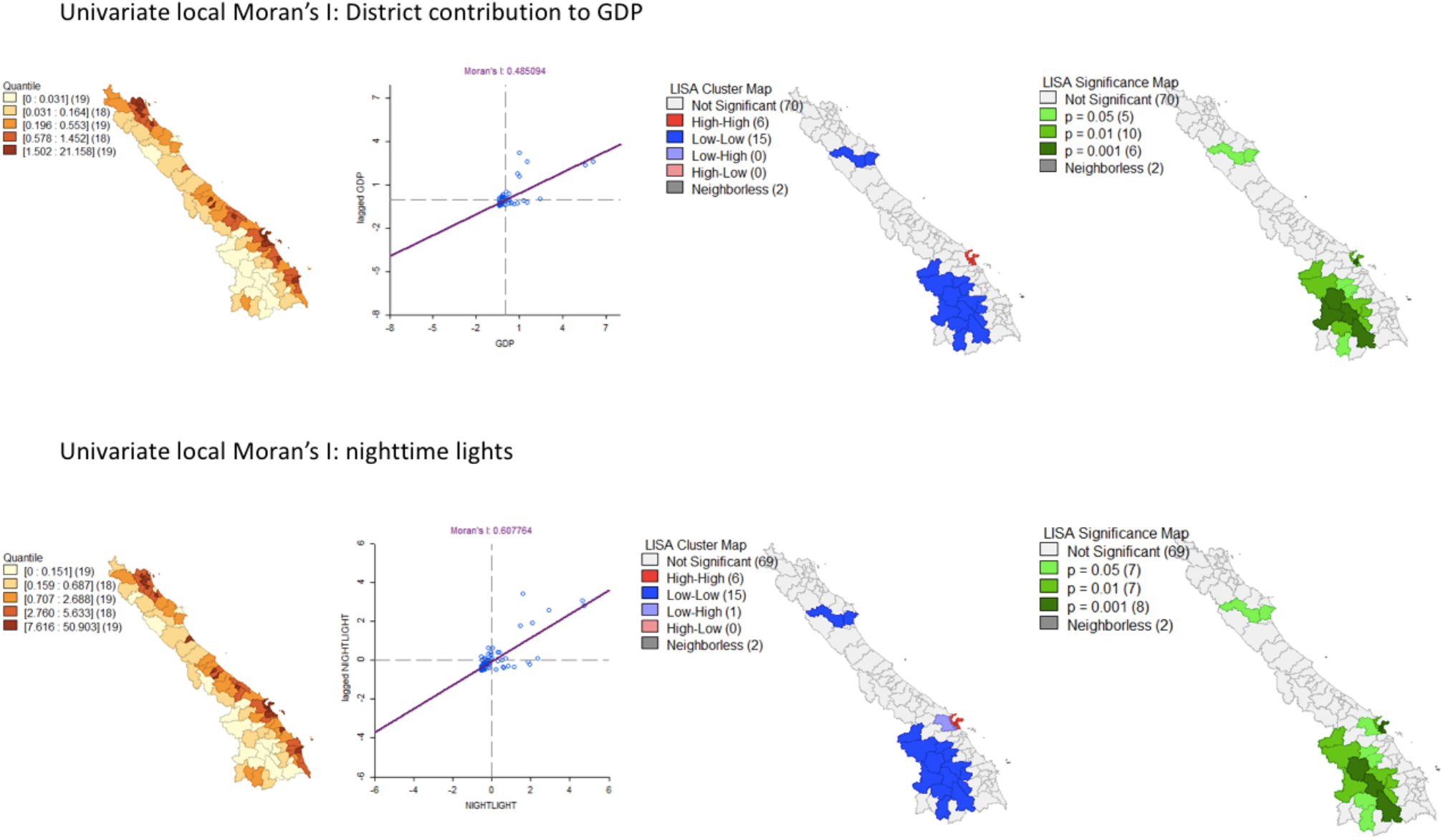
Mean coverage maps by district and univariate local Moran’s I of human development variables, district contribution to national GDP and nighttime lights, in CAL. Both display significant spatial autocorrelation (Moran’s I: 0.485, 0.608, respectively) and so spatially lagged variables were constructed.

**Figure 17.**
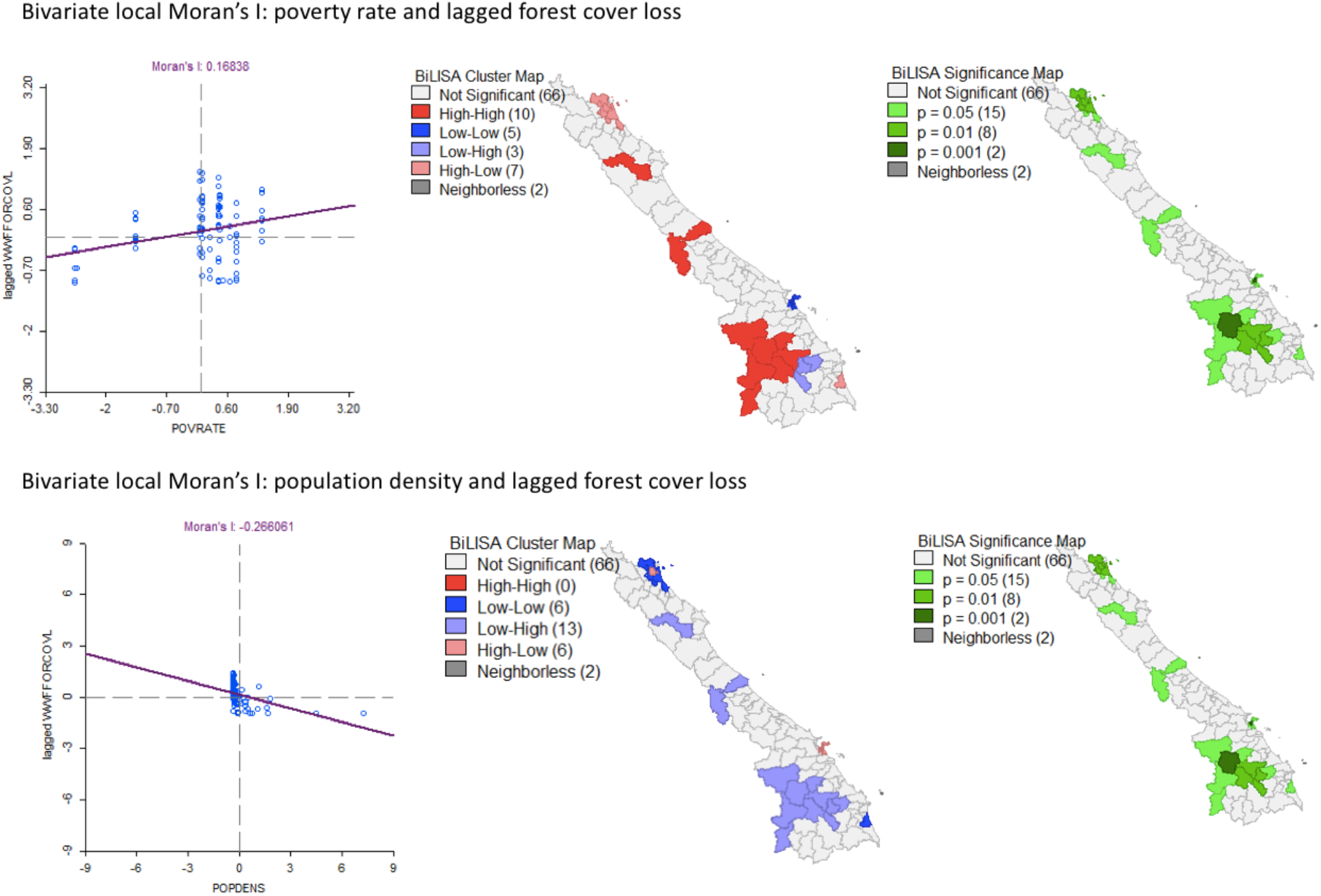
Bivariate local Moran’s I of poverty rate and population density relative to lagged forest cover loss (essentially, the “neighborhood” of forest cover loss), in CAL. While high poverty rates are spatially associated with forest cover loss (0.1684), population density is not (−0.2661). This provides some evidence that population pressures in the forest frontier areas may not be driving deforestation, but poverty and expansion of subsistence agriculture are, from a spatial standpoint, more likely determinants.

**Figure 18.**
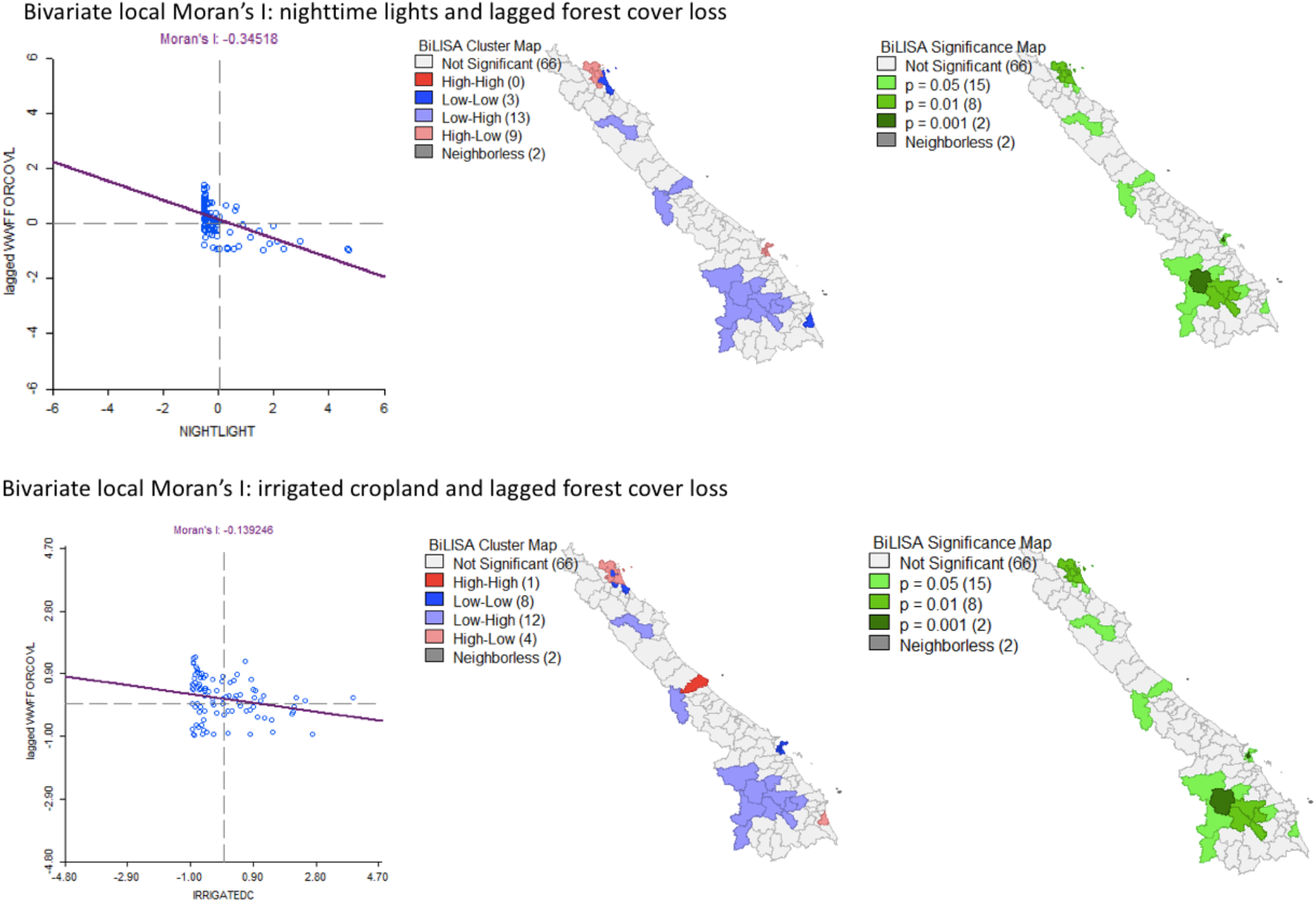
Bivariate local Moran’s I of nighttime lights and irrigated cropland relative to lagged forest cover loss (essentially, the “neighborhood” of forest cover loss), in CAL. Neither is spatially associated with forest cover loss (−0.3452, −0.1392, respectively). From Figure 8 and evidence from the univariate local Moran’s I in previous figures, the hotspot areas of deforestation are not spatially collocated with population-dense, heavily economic areas. This figure illustrates that they are neither potentially collocated with intense irrigated agriculture, suggesting that (and corroborating Figure 8) higher poverty rates and smaller scale subsistence agriculture may be driving deforestation at the immediate scale.

**Figure 19.**
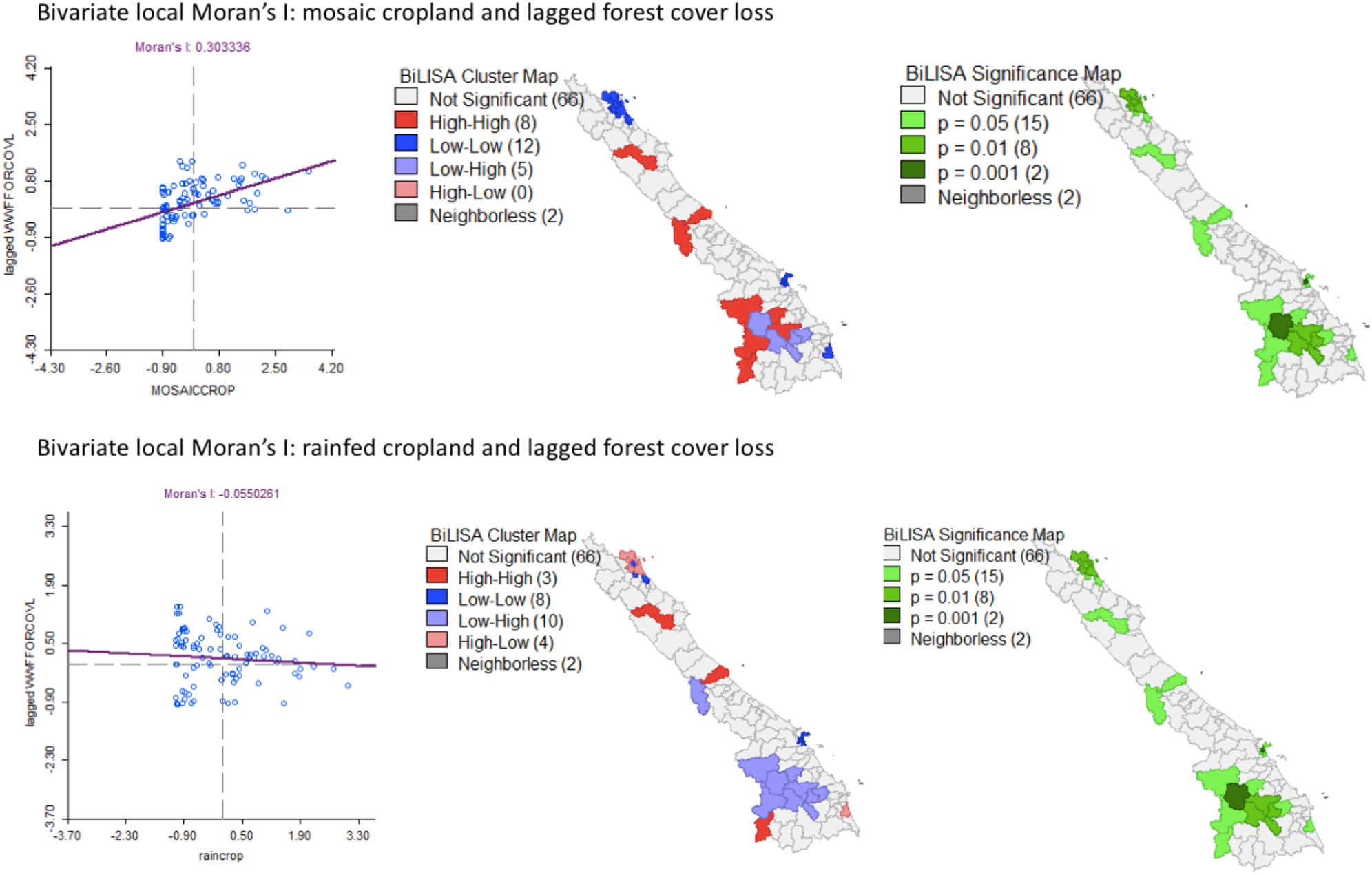
Bivariate local Moran’s I of mosaic cropland and rainfed cropland relative to lagged forest cover loss (essentially, the “neighborhood” of forest cover loss), in CAL. Mosaic cropland is spatially associated with forest cover loss (0.3033). Rainfed cropland is not spatially associated with forest cover loss but is less dispersed than irrigated cropland relative to forest cover loss in Figure 9 (−0.055 compared to −0.139).

Larger scale agriculture and other intense economic activities that are a known cause of deforestation in the landscape may thus be driven by conditions further afield, such as broader economic conditions as will be discussed further in the next section, and therefore this deforestation would be mediated by land tenure and property rights institutions. With government or other institutional support, property rights regimes could be adjusted to disincentivize deforestation.

#### 3.1.2 Underlying Causes Model

In this section we present the results from our assessment of underlying determinants of deforestation in CAL, macroeconomic trends linked to deforestation. First, Tables 4 and 5 are control specifications which highlight the relationships between biophysical control variables, cropland classifications, and tree cover loss in CAL (Table 10 also shows relationships between variables prior to accounting for spatial autocorrelation). Next, due to data inconsistencies in spatial and temporal variation in addition to high levels of collinearity in selected variables, we split similarly trending variables into separate specifications. To illustrate why this step was necessary, Table 3 presents a correlation matrix between prices and Vietnamese GDP. Not only are the selected prices and indices presented highly correlated with GDP, they are also correlated with each other. We expect some degree of this given the aggregation of dense time series to an annual time scale (where prices and indices fluctuate at extremely fine [daily] time scales) and lack of spatial variation. We also may expect that global prices and price indices will track, to some degree, on moderately similar trends at the annual scale given the interconnectedness of the global economy and commodity markets. However, the very high degree of collinearity is problematic for “kitchen sink” models that include all variables, hence our separated and restricted specifications.

The results of the Underlying Causes model are presented in Tables 6 and 7, with a combined restricted model in Table 9. Contrary to EKC theories, and in alignment with the literature, it appears that economic growth and increasing trade are associated with tree cover loss in CAL (Bhattari and Hammig, 2001). Table 6 M1 provides evidence that each percent of annual Vietnamese economic growth is associated with approximately 1.99 ha of tree cover loss in CAL and increasing trade may be associated with up to 20.90 ha of tree cover loss annually. Each percent of Chinese GDP growth is associated with a loss of 3.96 ha of forest cover, and similar increases in Chinese trade may be associated with 1.72 ha of tree cover loss in CAL annually. These variables maintain the direction of the effect (in terms of increasing tree cover loss) but lose their significance in Table 7. Rising trade in neighboring economies Laos and Myanmar (also collinear with Cambodia) are associated with tree cover loss, although these were inconsistent from Table 6 to Tables 7 and 8. India and Japan are also strong regional economies significantly associated with tree cover loss. The results are less clear for other regional and neighboring economies. Importantly, many of these neighboring and regional economies are interwoven and therefore endogenous; the result for India and Japan may actually be driven by China’s economic growth and its effect on the relationship between India and Japan’s economies and deforestation.

Of commodity prices and price indices assessed, rubber and hardwood price present the greatest magnitude and consistency in their effect on tree cover loss. Increases in rubber price are consistently associated with increasing tree cover loss, both within the same year and within a one-year time lag. A one-unit increase in rubber price annually can result in a loss of between 2.7 and 10.5 ha of forest cover, and such an increase can result in 7.9 ha of loss in the following year. Hardwood price had a large relative magnitude of effect, as a one-unit increase in the value of the index was associated with a 49.79 ha loss in forest cover in Table 6. Increases in metals price were associated with increasing tree cover loss in specifications where it was statistically significant, but its significance was inconsistent across specifications. Food commodity prices such as cereals and rice were also less consistent, although the tea-coffee-cocoa index, like metals prices, maintained the direction of effect (increasing tree cover loss) in all specifications where it was statistically significant. For rice prices, the relationship changes between within-year and the temporally lagged rice price in Table 6.

#### 3.1.3 Immediate Causes Model

The results of the Immediate Causes model are presented in Table 8, with a combined restricted model in Table 9. Both fixed-effects and standard spatial regression are included in the table. Population density is not significant in any of these specifications, likely due to lack of spatial collocation with forest cover. Poverty rate is consistently associated with tree cover loss, while urban area is consistently (though insignificantly) associated with reduction in tree cover loss. This may be due to a combination of lack of spatial collocation, like the population density variable, and increased off-farm wage and employment opportunities in urban areas. That said, while significant economic activity is not spatially associated with tree cover loss, those districts contributing more to overall Vietnamese GDP appear to have greater tree cover loss than those contributing less to overall GDP. This contribution to GDP could be in agriculture, but this result highlights a need for further data collection and research on each districts’ primary economic activities that contribute to GDP, with spatial and temporal variation. Importantly, districts with higher secondary school graduation rates demonstrate reduced tree cover loss, and this trend was consistent and significant across all specifications. Table 8 M1 indicates that increasing the rate of secondary school graduation by 1% can reduce tree cover loss by as much as 12.7 ha. Of agricultural land types, rainfed agriculture and mosaic cropland appear to have the greatest effect on tree cover loss. While Table 5 M3 shows that irrigated cropland was significantly associated with tree cover loss, this was not found in any specification that included biophysical control variables.

#### 3.1.4 District and Provincial Sensitivity

Based on these results, particularly the combined restricted model in Table 9, we ran district and province-specific regressions to highlight the relative effects of key variables, and then created “sensitivity maps” that visualize which districts and provinces of CAL are the most sensitive to the effects of deforestation determinants by magnitude. Tables 11 and 12 present the results of those factor variable (district interaction) models and Figures 20 and 21 present the mapped visualizations of these regressions. The results indicate that the districts with the most sensitivity to fluctuations in underlying economic conditions are located in the provinces of Quảng Bình, Quảng Tri, and Thù’a Thiên Huế. Returning to Figure 2, these are also provinces with a large number of frontier forest districts, where the areas of high forest cover begin to meet the most heavily economic and population-dense coastal areas. As such, both high tree cover loss and high sensitivity to economic conditions exist in districts of those CAL provinces. Importantly, Figure 20 (bottom right map) illustrates that a number of districts with high tree cover loss are also highly sensitive to improvement in immediate economic conditions such as educational attainment by secondary school graduation rate. The districts of Bắc Trà My, Trà Bồng, Cam Lộ, Triệu Phong, Hương Trà, and Đa Krông are strategically both higher tree cover loss and sensitive to this immediate driver.

**Figure 20.**
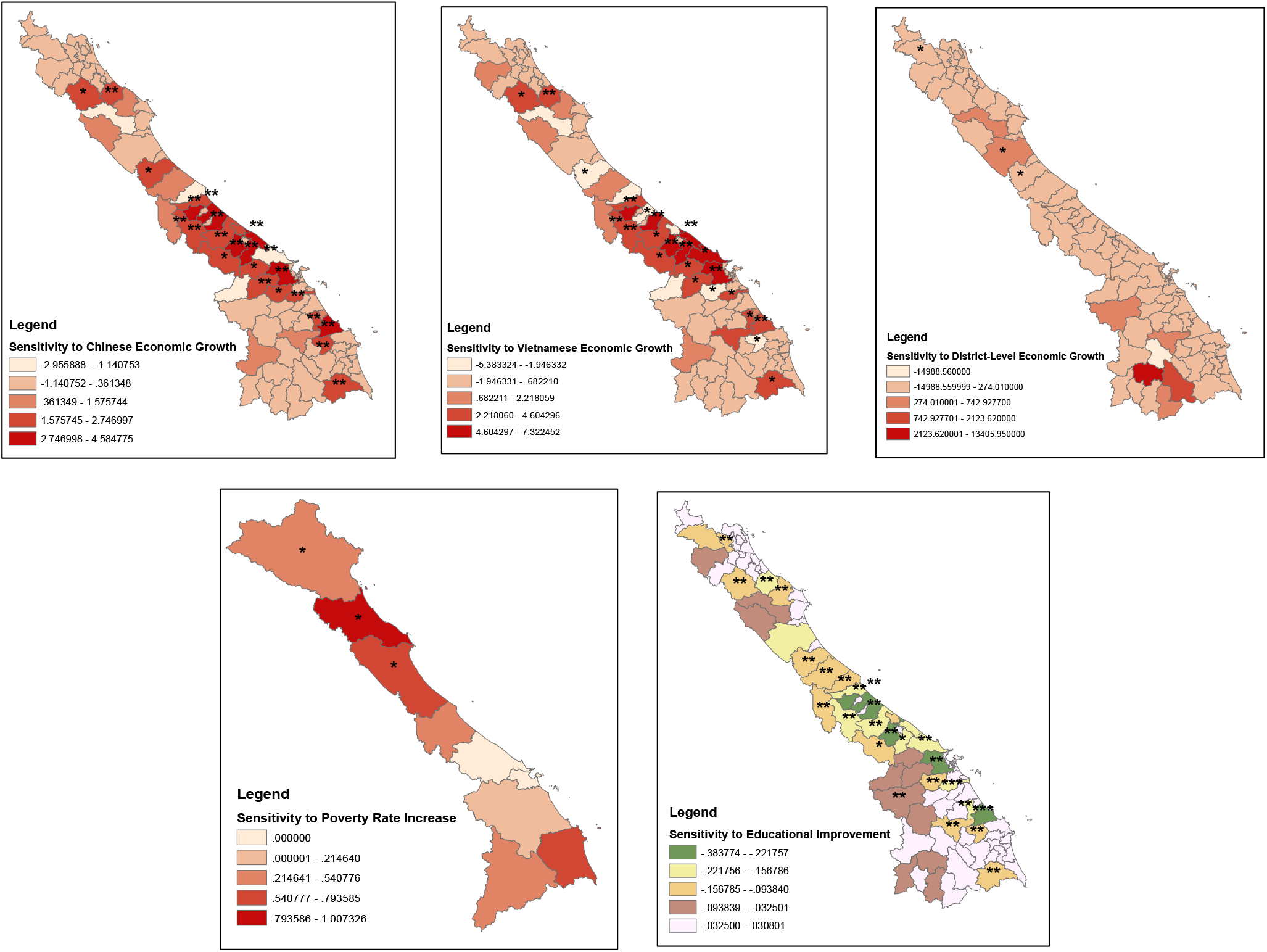
Sensitivity of deforestation to explanatory variables, based on district-level regression coefficients from Tables 11 and 12. *** p<0.01, **p<0.05, *p<0.1. Generally, the districts most sensitive to economic conditions are located in the provinces of Quảng Bình, Quảng Trị, and Thừa Thiên Huế.

**Figure 21.**
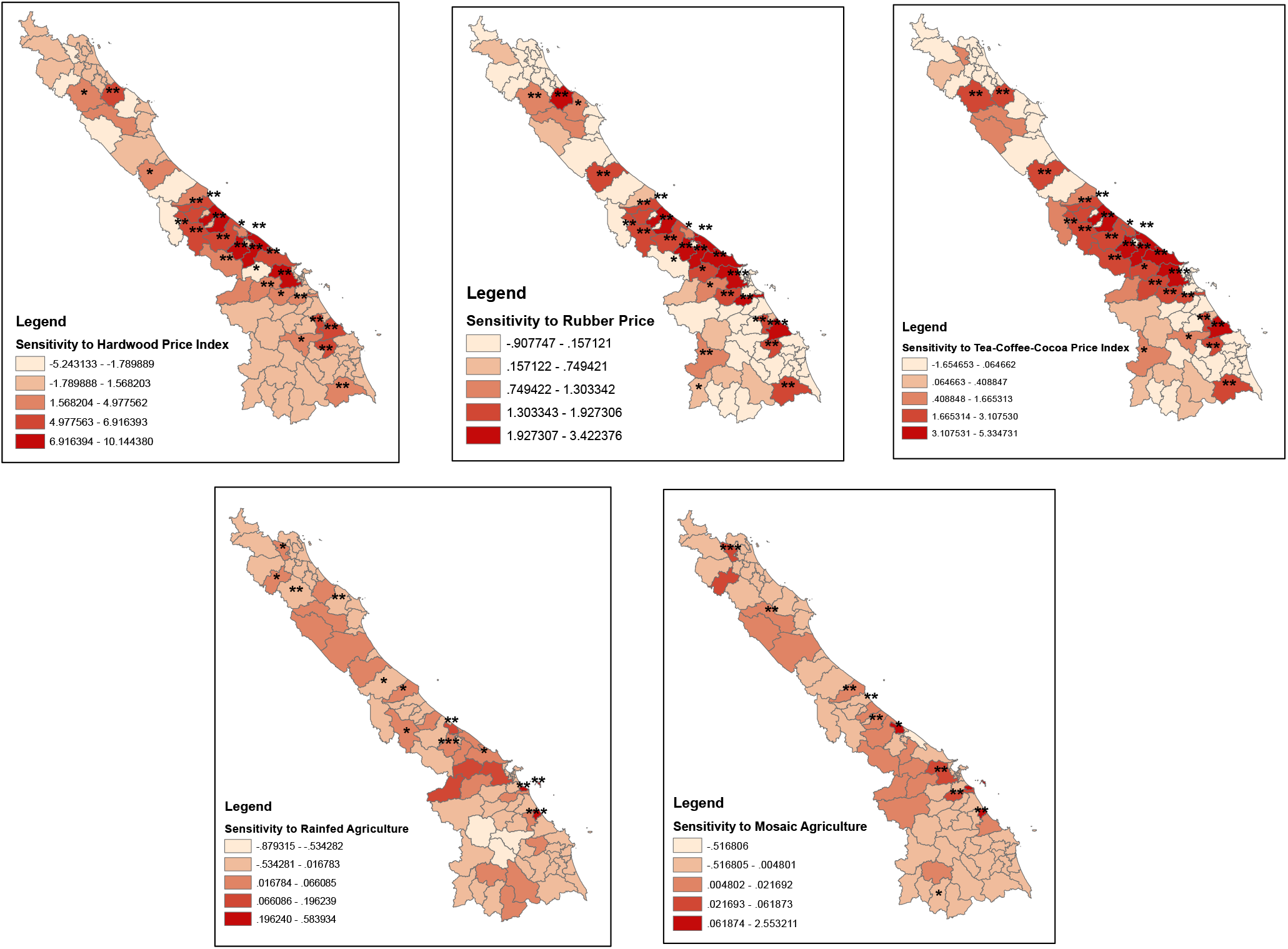
Sensitivity of deforestation to explanatory variables, based on district-level regression coefficients from Tables 11 and 12. *** p<0.01, **p<0.05, *p<0.1. Generally, the districts most sensitive to economic conditions are located in the provinces of Quảng Bình, Quảng Trị, and Thừa Thiên Huế.

We used the results of these Underlying and Immediate Determinants models, and the district-specific sensitivity specifications, first to create a “deforestation risk map,” and then to select models for Phase 2, using predictive margins for scenarios and target feasibility analysis. The deforestation risk map in Figure 22 is a weighted overlay of the following reclassified rasters: tree cover loss, which is given 50% weight; inverse distance weight (IDW)-interpolated rasters of underlying economic conditions from Table 9 M4 regressions and their subsequent sensitivity maps, given 10% weight; and immediate socioeconomic conditions variables from Table 9 M4 regressions, given 5-10% weight depending on magnitude and significance. The reclassifications were based on a 1-9 scale, separating the values of the sensitivity regression coefficients and interpolation values into integer values 1-9 in equal breaks. The layers with these values are then weighted as previously discussed, with actual historical tree cover loss as primary (50%), and overlaid using ArcGIS’s Weighted Overlay tool. It is evident from this mapping that the districts in the province of Quảng Trị, including Hướng Hóa, Đa Krông, and Cam Lô, and other districts such as A Lưới, Minh Hóa, Hương Khê, and Đắk Glei display notable deforestation risk based on history of tree cover loss, sensitivity to various underlying economic conditions, and recent values for immediate socioeconomic conditions.

**Figure 22.**
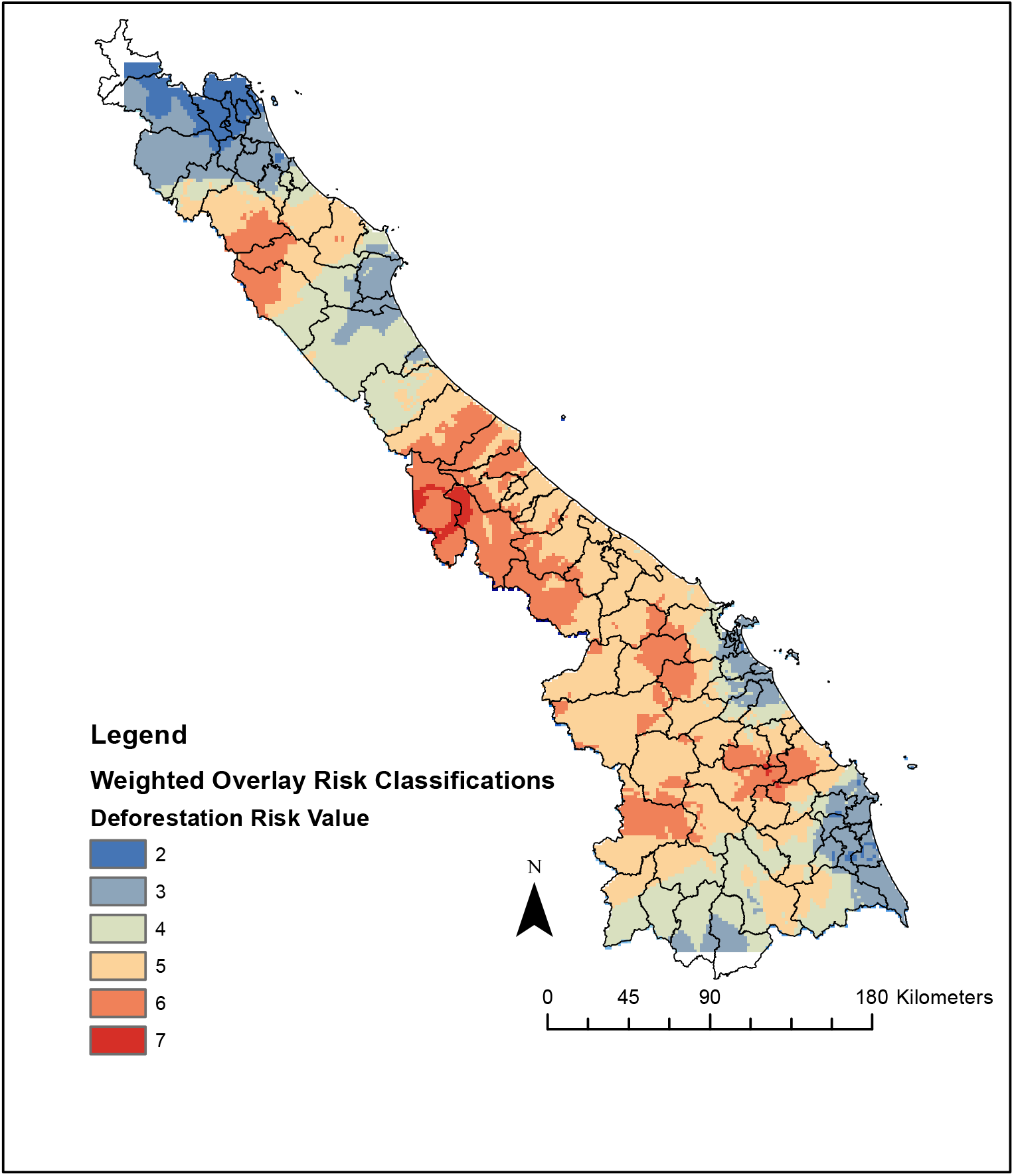
Deforestation “risk map” based on a weighted overlay of historical forest cover loss (50% weight), reclassified inverse distance weight (IDW)-interpolated rasters of underlying economic conditions from Table 9 M4 regressions and their subsequent sensitivity maps (10% weight), and immediate socioeconomic conditions variables from Table 9 M4 regressions (5-10% weight depending on magnitude and significance). Districts in the province of Quảng Trị, including Hướng Hóa, Đa Krông, and Cam Lô, and other districts such as A Lu’ϳ’i, Minh Hóa, Huong Khê, and Đắk Glei display notable deforestation risk based on history of forest cover loss, sensitivity to various underlying economic conditions, and recent values for immediate socioeconomic conditions. Higher values (red) indicate greater “deforestation risk.” The value “1” is not included as this contained “no data” values (white, outside of the extent of the forest cover loss raster).

Next, Table 13 illustrates the results of predictive margins for scenarios for tree cover loss in four areas: Chinese economic growth, rubber price fluctuations, poverty rate changes, and educational attainment. These are to be cautiously interpreted as the models used had already low R^2^ values for any kind of predictive modeling. These methods were also used to fix the most recent economic conditions and thus estimate approximate expected minimum deforestation rate assuming that these economic conditions continue. This can assist WWF in determining the feasibility of targets for reducing deforestation. These results are discussed further in the following section, Phase 2.

### 3.2 Scenarios of economic and socioeconomic conditions

Following the Phase 1 modeling, we selected Chinese economic growth, rubber price fluctuation, poverty reduction, and secondary school graduation for use with predictive margins. We estimated the models presented in Table 13. The specific model used is indicated underneath the equation in each cell, along with the fixed values to predict the possible effect on deforestation for each scenario. Fixed values were selected using the minimum, maximum, and median values for each variable. We followed the most successful model utilizing each variable, prioritizing R^2^ values. *X* is a vector of non-interest variables (the remainder of variables run normally in the regression) for a log-log spatial OLS. *ε_dpt_* is the error term for district *d*, province %, and at time *t*.

Table 13 provides a simple classification of these results, based on whether the coefficient for that variable’s effect on tree cover loss increased, decreased, or remained the same from the coefficient for that variable in the original model (indicated under the equation in each Table 13 cell). **Green** indicates that the given scenario does **is not associated with tree cover loss. Red** indicates that the given scenario **is associated with tree cover loss. Yellow** indicates that a significant relationship could not be reliably determined with a P-value under 0.10 using predictive margins. The results are in line with the regression results overall, with the exception of increase in rubber price. Surprisingly, fixing the rubber price at its maximum observed value in the dataset reduced the effect of rubber price on tree cover loss, contrary to the results of the other regression models. We recommend further research to explore this inconsistency.

We estimated that maintained growth of Chinese GDP may predict increased deforestation, although we were unable to estimate a reliable prediction (p>0.10) for economic contraction or economic boom. In contrast to the previous econometric outputs, we found that increase in rubber price may predict reduced deforestation, but this predictive margin was of borderline significance (p=0.1000). Next, a worsening of poverty rate was estimated to predict worsening deforestation, while increase in secondary school graduation rates beyond current levels was estimated to predict reduced deforestation.

#### 3.2.1 Target feasibility analysis

We used predictive margins to further estimate minimum overall tree cover loss across CAL, given the most recent observed values for a number of economic variables. For strategic planning purposes, this can be useful for determining expected deforestation in the absence of interventions. We estimated

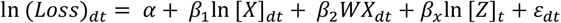

based on Tables 9, 10, and 13, where X is a vector of the same controls included in those models. We selected key economic variables also identified in those models and fixed their values in the vector Z, which included:

- China GDP (16806.742 [PPP] 2017; log transformed) =9.65057
- Vietnam GDP (6775.83219 [PPP] 2017; log transformed) =8.747604
- Rubber price (90.790599 [USD per metric tonne] 2016; log transformed) =4.310578
- Hardwood price index (124.791 [index 2005=100] 2016; log transformed) =4.82664
- Metals price index (126.574 [index 2005=100] 2016; log transformed) = 4.940827
- Cereals price index (148.98 [index 2005=100] 2016; log transformed) =5.003812
- Tea-coffee-cocoa index (172.587 [index 2005=100] 2016; log transformed) = 5.150901
- Secondary school graduation (95.8 [% average] 2017; log transformed) =4.5

Our resulting predictive margin given these fixed values was 7.547895** (p=0.004) with an original model R^2^ of 0.652, and our 95% confidence interval predicted a range of [2.591826: 12.50396]. Using the conservative end of the range, current market conditions would predict minimum tree cover loss of approximately 2.59% with 95% confidence. This would not be uniform across CAL, and as Figures 20 and 21 illustrate, districts have varying degrees of sensitivity to the economic conditions included in this model. There are also certainly other non-economic conditions, such as changes in local or national forest policy or governance that could greatly impact this result. Further, variation in institutions such as land tenure and property rights can greatly affect land use change in CAL districts.

## 4. Discussion

In this section, we summarize overall findings from our exploration of socioeconomic determinants of deforestation across Vietnam’s Central Annamites Landscape, and tie together the distinct-specific and spatial elements of this pilot study.

First, the data availability assessment is a fundamental component of our study (Appendix II), as it has shaped variables used to perform these analyses. With more temporal and spatial variation in just a few of the variables, much of the constraints resulting in coarse and correlative analysis, required by collinearity, could be avoided. An important next step for future research will be determining the approach for acquiring data that is online but currently inaccessible, such as GSOV household living conditions survey data or VNForest data available for visualization at a fine scale online.

For underlying macroeconomic determinants of deforestation in CAL, we found evidence to suggest that Vietnamese GDP growth and that of neighboring and regional economies, such as China, have been associated with increased tree cover loss in CAL. Overall economic and development pressures are thus underlying drivers of activities that result in deforestation; we did not find evidence to support a deforestation Environmental Kuznets Curve in CAL, in alignment with the Bhattari and Hammig’s findings (2001). While we found through spatial analysis that these economic pressures, including those from population-dense and urban areas, are not spatially collocated with the areas of highest tree cover loss, we did find that districts in the provinces of Quảng Bình, Quảng Trị, and Thừa Thiên Huế are sensitive to fluctuations in underlying economic determinants of deforestation, and these districts are strategically located along the transition from urban/economic coastal areas to the western forest frontier (Figures 1, 20, 21).

Our findings suggest that components of contribution to GDP and land conversion taking place within these districts needs to be broken down further to better understand the implications for deforestation. The results from district individual contribution to GDP were not in alignment with those from districts with higher urban area. District contribution to GDP was associated with increased tree cover loss in several specifications, while higher urban area was consistently associated with reduced tree cover loss. This is potentially due to the increased off-farm wages and employment opportunities in urban areas but indicates a need to understand which specific contributions each district makes to national GDP. Future research can thus pinpoint which specific activities within each district are contributing to the tree cover loss associated with that district’s contribution to GDP. Considering agricultural contributions to GDP, this could help to further isolate specific commodities driving land use change and tree cover loss (Table 6 M1 and M2; Figure 17, Figure 18).

Next, we used commodity prices that provide an underlying signal for land use change, and therefore can proxy for land use change as an underlying driver of deforestation. Rubber, and hardwood prices had the most consistent and greatest magnitude of effect on tree cover loss, with indices for tea-coffee-cocoa and metals prices also associated with increased tree cover loss where significant. Food commodities such as rice and cereals were inconsistent. This aligns with findings of Cochard et al. (2013) and provides evidence to support current conservation efforts focusing on rubber, acacia, and artisanal mining. However, our finding of spatial collocation of deforestation with poverty and low population density, combined with the possible benefits of improved educational attainment across spatial analysis, economic modeling, and scenario analysis, indicates that immediate socioeconomic factors cannot be forgotten. Returning to the Angelsen and Kaimowitiz (1999) conceptual framework, the agent decision-making parameters are shaped by immediate socioeconomic conditions. In the simple scenario modeling, we found some evidence that lower poverty rates may predict more deforestation, while higher education attainment may predict reduced deforestation. This was also supported by the results of the district-specific specifications in Tables 11 and 12, and mapping in Figures 20 and 21. Deforestation particularly in the districts of Bắc Trà My, Trà Bồng, Cam Lộ, Triệu Phong, Hương Trà, and Đa Krông could be reduced through targeted efforts to improve immediate socioeconomic conditions. This evidence thus supports programs that encourage off-farm, off-plantation, and off-mine employment and educational opportunity.

Finally, the feasibility of conservation targets and associated programs can be determined in the context of continuing economic determinants of deforestation. We cautiously predict annual deforestation of approximately 2.59% in CAL, with spatial inconsistency, given 2017 economic conditions. This figure is quite high, but these conditions undoubtedly fluctuate, and the actual deforestation rate is mediated by property rights regimes, land tenure and concessions, and the local and national institutional and regulatory environment which are not included in this model. However, we emphasize that economic conditions, particularly regional and global-scale, underlying economic forces, will continue to drive deforestation at a fairly high rate. Strategic planning for forest conservation will take this into account in setting targets, prioritizing spatial locations, allocating program and project funding to areas of greater risk. Overall, Figure 1 illustrates that the greatest average tree cover loss by magnitude in CAL is in Quàng Nam province, but Figure 22 provides evidence that the forests in Quảng Trị province are more at risk due to sensitivity to economic conditions, both underlying and immediate. Quảng Trị districts, particularly Hướng Hóa, Đa Krông, and Cam Lộ, in addition to Quảng Bình and Thừa Thiên Huế districts A Lưới, Minh Hóa, Hương Khê, and Đắk Glei, display notable deforestation risk which should be incorporated into strategic planning.

## 5. Conclusion

In conclusion, this study has used tiered spatial regression analysis to correlatively identify which commodities, economic development activities, and social conditions have historically had the greatest effect on forest cover by magnitude in CAL. Based on first phase results, we selected a subset of these determinants for scenario modelling to predict possible deforestation outcomes given certain economic scenarios.

The results, among others, indicate that terms of spatial collocation, high poverty rates and smaller scale agricultural land conversion are key immediate determinants of deforestation. That said, further research is required to specifically break down the types of agricultural land use that are responsible. In the absence of land use data with spatial and temporal variation, we break down these land use types using the proxy of commodity prices. Price or index value increases for rubber, hardwoods, metals, and a tea-coffee-cocoa index are consistently associated with tree cover loss, ranging from approximately 2 to 16 ha of tree cover loss annually. This therefore provides evidence to support programs targeting rubber, acacia harvesting, artisanal mining, and land conversion for cash crop plantations. Education is also key immediate socioeconomic factor, as poverty rate is consistently associated with tree cover loss and increased educational attainment is consistently associated with reduced tree cover loss, at approximately 12 ha per percentage increase in secondary school graduation. On the macro level, economic growth in China and Vietnam are correlatively associated with tree cover loss, as are rising trade in Myanmar and Laos. This study provides a methodological contribution to the current academic literature identifying socioeconomic dimensions of deforestation through spatial econometric analysis and scenario modelling at the landscape level. For practitioner work, we hope that this pilot study provides a tool for strategic planning of conservation interventions in light of economic conditions and factors, not only through the empirical analyses conducted but also through our systematic assessment of currently available data for understanding these issues. As a pilot, we hope to highlight areas of need for data collection and future research.

Based on the outcome of the data availability assessment and spatial and quantitative results of this baseline study, we recommend future research focusing on immediate drivers of deforestation (micro/socioeconomic determinants) should utilize currently available data by focusing on small scale target areas within landscapes. This will necessitate working with GSOV to acquire the household-level data from the household living conditions surveys that is not available online, or collecting new data by administering a small household living conditions survey focusing on off-farm wage opportunities and opportunities outside of timber, rubber, and other heavily-deforesting harvesting. With the baseline established in this study, future work can focus on using quasi-experimental methods such as border or regression discontinuity based on spatial policy differences or temporal “shocks.” Instrumental variables could also be explored. Lao data could be added for CAL, enabling border discontinuity design for economic and conservation programs on the Vietnamese side. Further, the economic growth contraction in Cambodia in 2009 could be a source of “shock” for a natural experiment, but this needs to be explored.

A key missing element in this analysis is local-level governance, which can vary across provincial and district levels. Following Khuc et al. (2018), a provincial level governance variable could be included in future studies. Additionally, property rights regimes and land tenure systems must be incorporated into this analysis as well to account for institutions which are immediate determinants of deforestion (Angelsen and Kaimowitz, 1999; Angelsen, 2010; Agarwal et al., 2008).

We also note some key strategy considerations for organizations involved in integrated conservation-development in Vietnam. Economic conditions and their effect on deforestation varies across districts in the CAL, and continuing rates of deforestation driven by these factors must be taken into account throughout project and program planning and implementation. Evidence supports conservation efforts focusing on rubber, acacia/hardwood logging, and artisanal mining, but socioeconomic conditions of agricultural communities near forests cannot be forgotten.

## 7. Acknowledgements

This pilot study was conducted and working paper drafted initially as a World Wide Fund for Nature (WWF) – Greater Mekong report in 2018-19. The authors are grateful to WWF Vietnam, and to Benjamin Rawson, Thibault Ledecq and Tam Le Viet for data collection support, review, and comments. This project received funding from WWF Greater Mekong and WWF Switzerland.

## 8. Author contributions

KPB: conceptualization, data curation, data management, methodology, formal analysis, writing original draft, writing review and editing, visualization

SZ: conceptualization, project management, methodology, writing review and editing

ACS: conceptualization, methodology, writing review

## Appendix I: Summary statistics

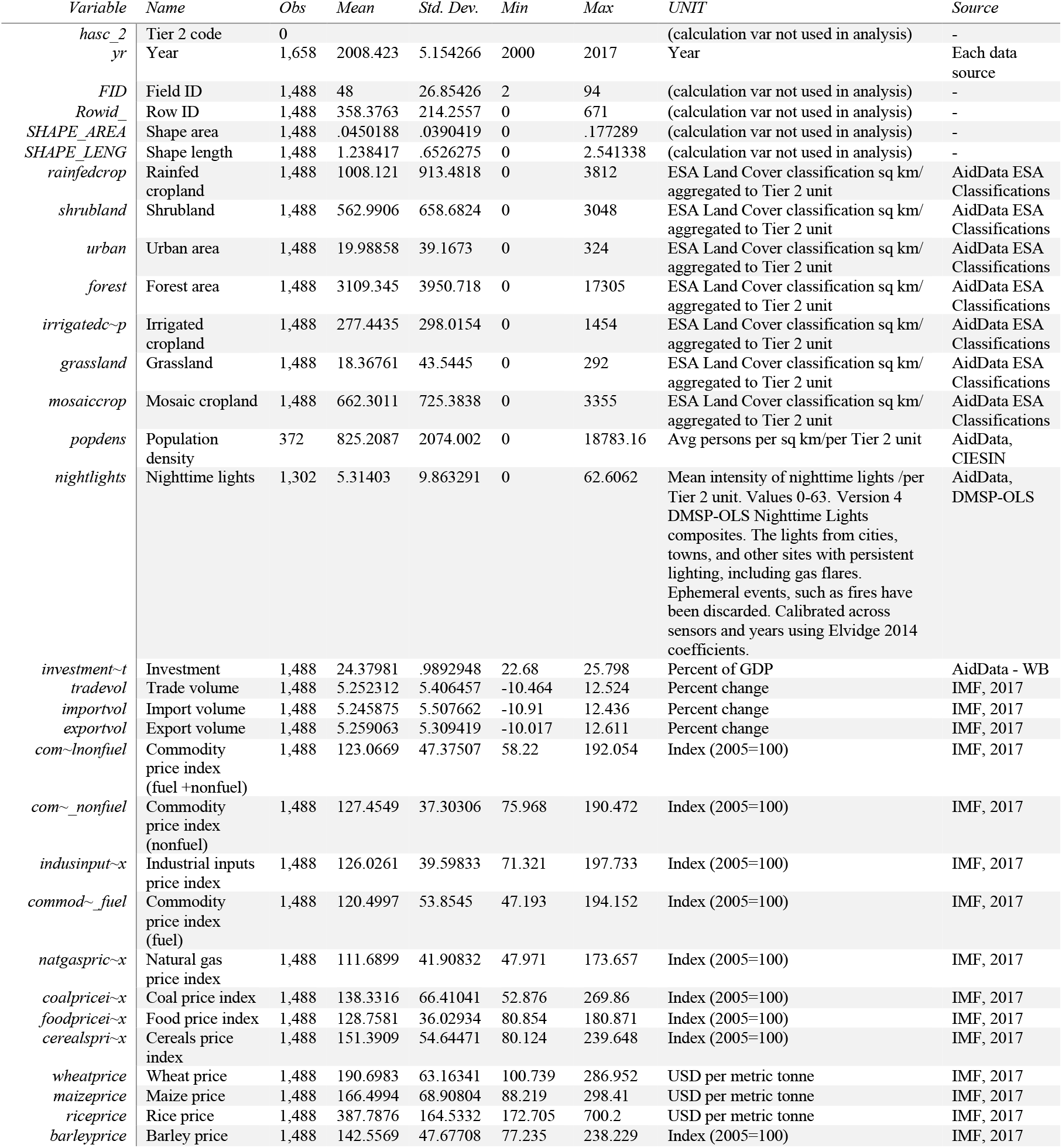
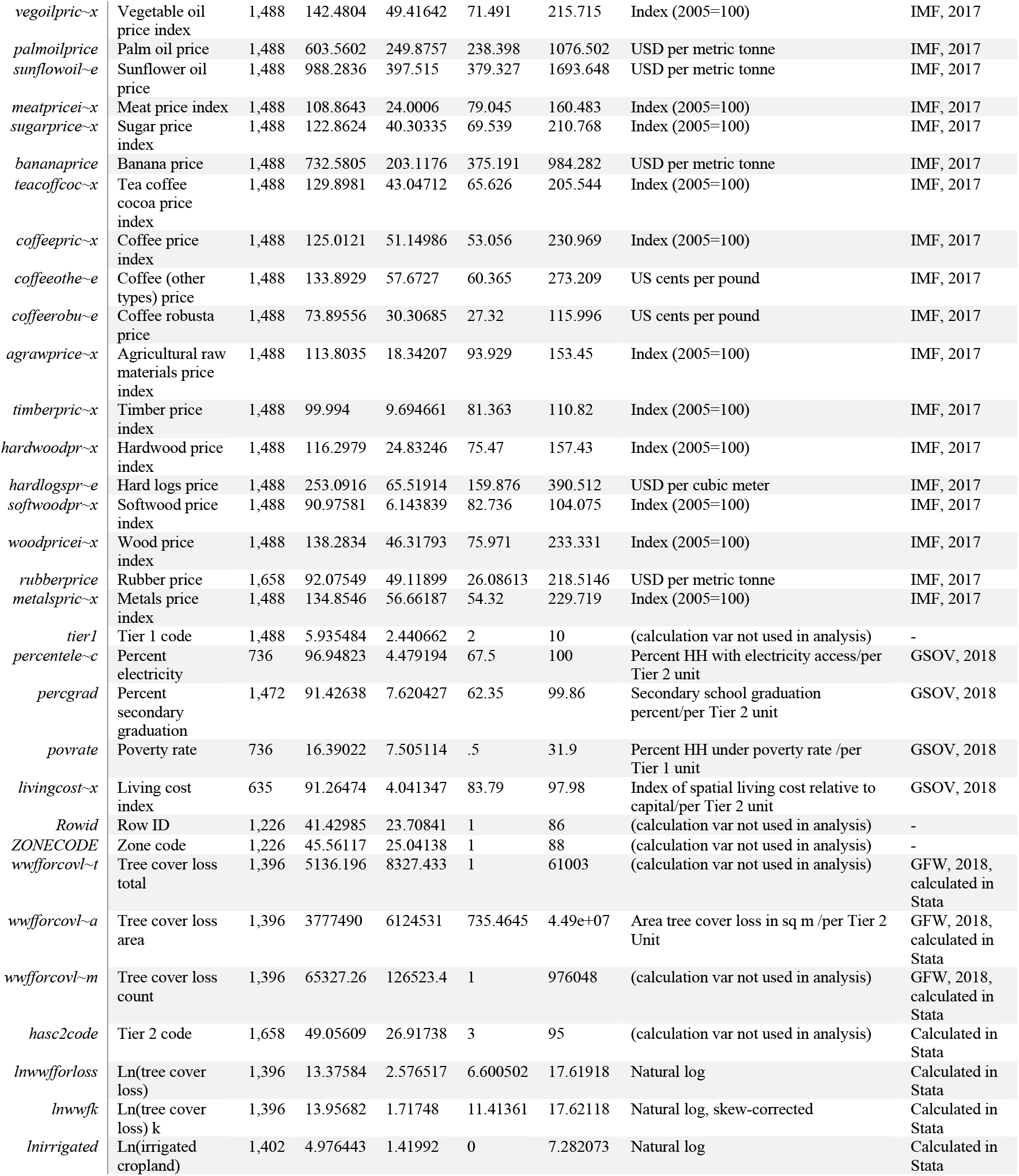
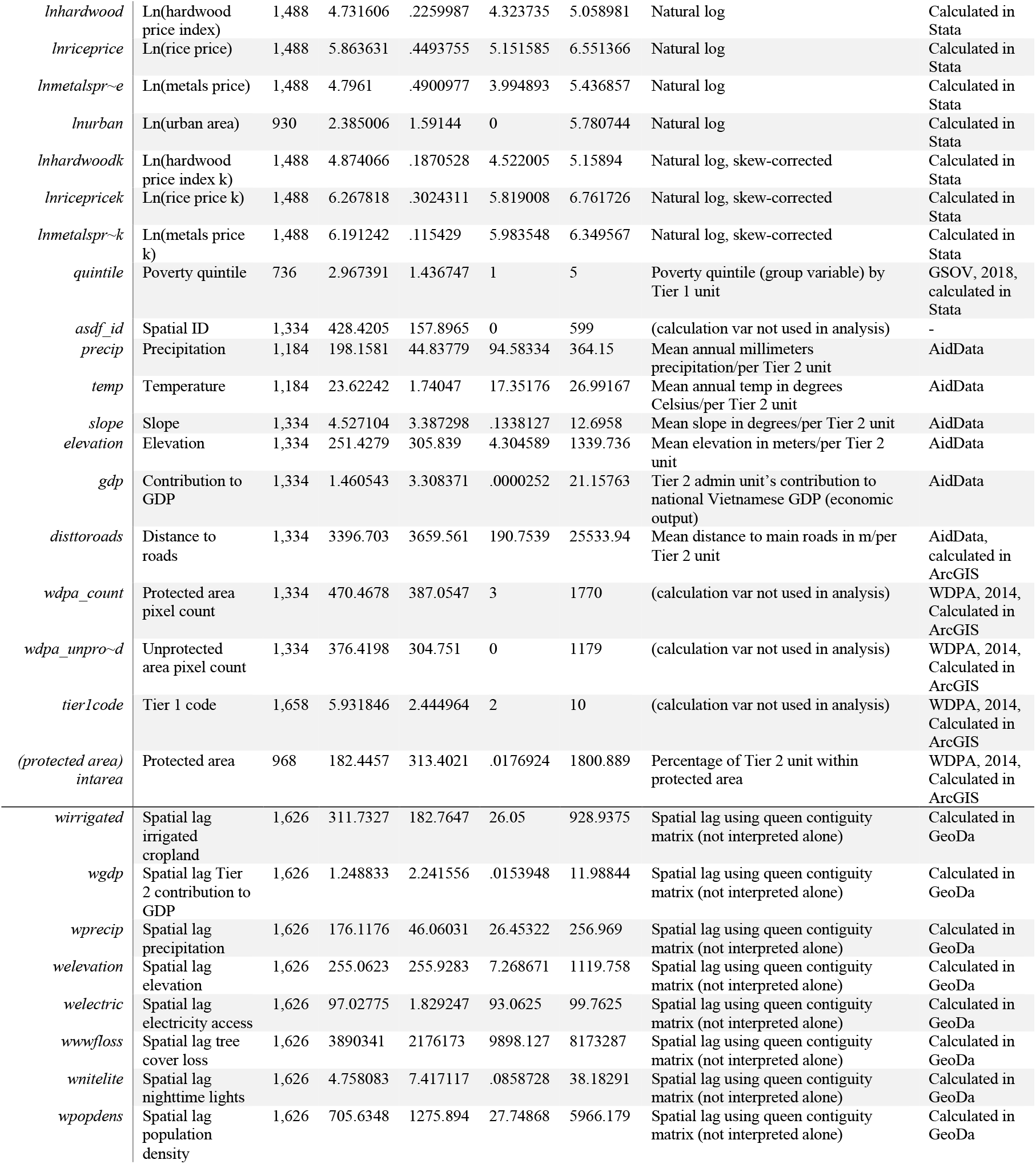
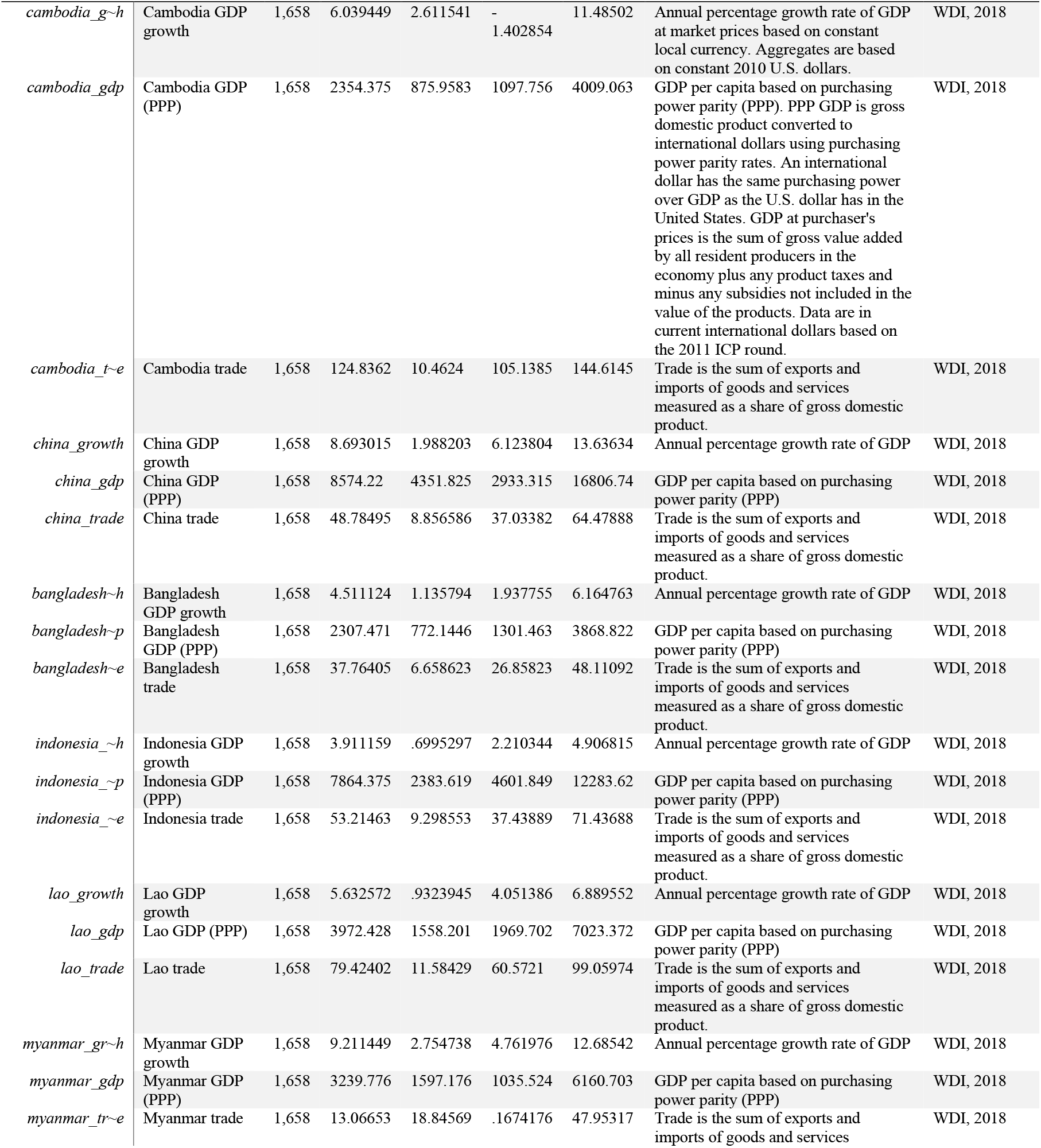
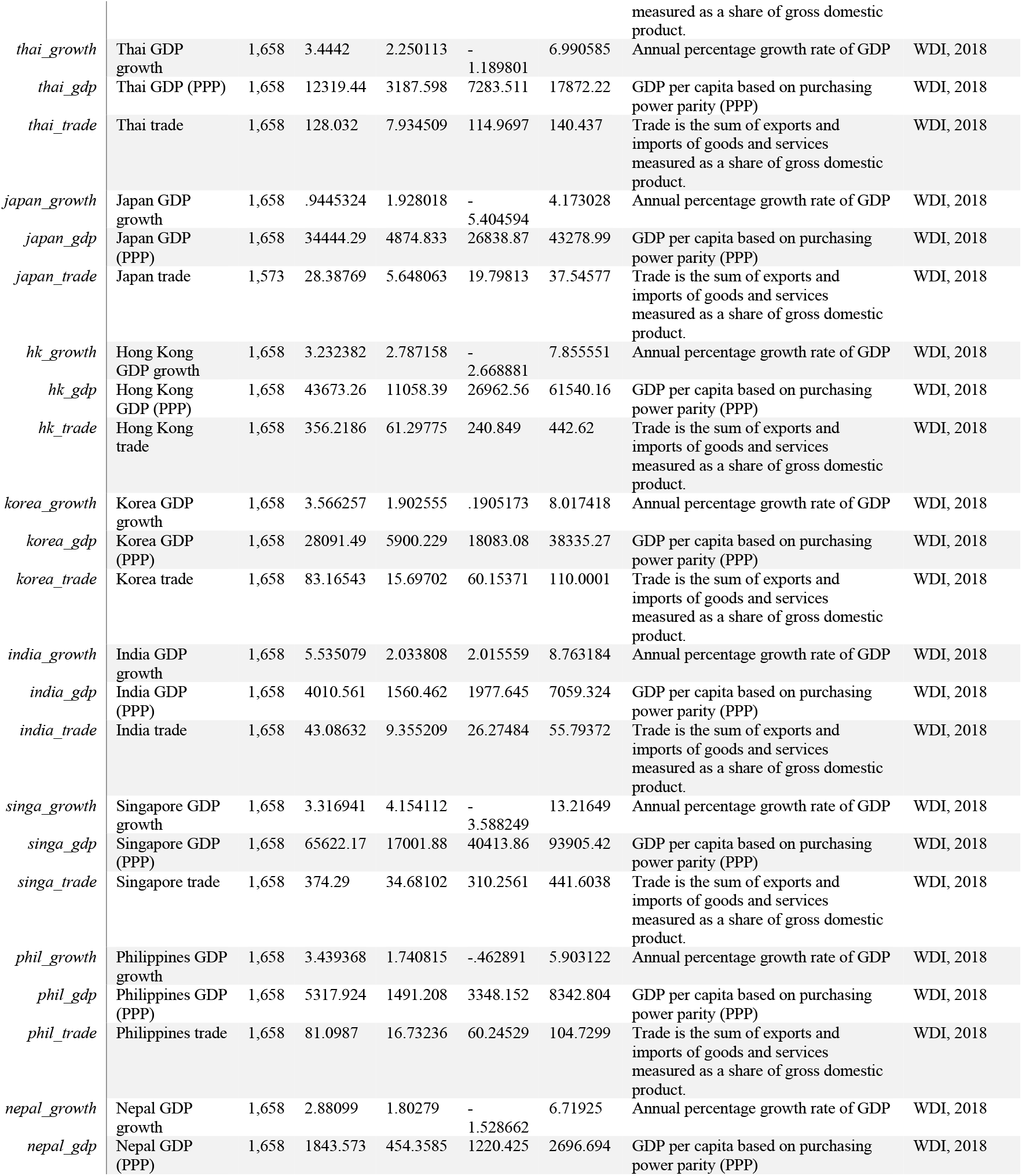
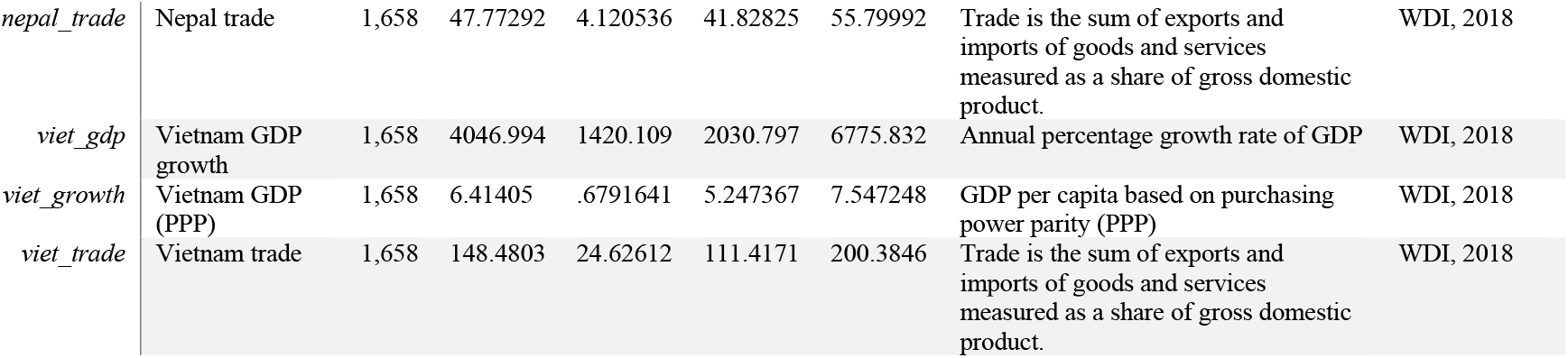

## Appendix II. Data availability assessment

Reliable methods for estimation using spatial panel econometrics require data from multiple specific administrative units or spatial subareas, which can be difficult to obtain in the absence of annual, or at least periodic, raster data or consistent government data collection. We sought data at the Tier 2 administrative level in order to establish sufficient observations in a panel for statistical robustness. In addition to the need for multiple spatial units, these methods become more reliable with more periods of time series. We thus sought data for the time period 2000-2017, with data only available in waves (e.g., 2005, 2010, 2015) imputable for the entire period. In this regard, we summarize our findings of key areas of missing, incomplete, or available but inaccessible data at the landscape level with the following:

- **Socioeconomic household survey data** available in a usable **disaggregated** format online

- Particularly data for off-farm wages and poverty rates at the spatial and temporal scale required

■ GSOV data from household living conditions surveys is available in aggregated format for online download, however the disaggregated data is not available for download. The aggregated versions usually have one but not both of the spatial and temporal variation necessary. For example, we use poverty rate as a group categorical variable at the provincial level.
- Commodity-specific spatial **land use area with multiple temporal waves**

- E.g., raster image of rice agricultural area 2010 and 2015; raster image of timber production area 2010 and 2015

■ FORMIS data (from the Forestry Data Sharing System) is currently an excellent resource for fine-scale visualization, but the Data Catalogue is still under construction. Due to a technical problem on the website, were also unable to download the statistical reports at the district level for forest function sub-class. Therefore, while the online visualization of land use categories (Forest function sub-class: Watershed, protection forest for tide shielding, wind and sand shielding forest, environmental protection, national park, nature reserve, scientific research, historical and landscape area, big timber production, small timber production, bamboo production, other production) is useful, the data downloads are unusable for the purposes of this study. Furthermore, even if available, these would only be available for 2016 and 2017. The full raster or vector datasets would need to be made available for even the one-time-period use. However, this could still be useful for a much finer-scaled study, rather than at landscape level.
■ Product GFSAD30SEACE (Southeast Asia) in the Global Croplands project from NASA’s Land Processes Distributed Active Archive Center has only three classifications: water, non-cropland, and cropland.
- This would enable calculation of land conversion rate and comparison to price as a signal
- **District level economic activity data** including import/export volume, manufacturing/industrial production, local (Tier 2) commodity production and prices with **multiple temporal waves**

- GSOV data for this is available but usually either at district level OR over time, not both (over time is usually aggregated to country or provincial level); in this study we have extracted variables that do have both, however these were limited and constrained our analysis.

As a pilot study, we took a broad approach in secondary data collection and assessment of data availability. While we took care to identify and incorporate necessary control and explanatory variables, we used acceptable proxies and linear imputation when key variables were unavailable for the desired time period or entirely unavailable. Table 1 details the variable types used and theory backing those variable types in the literature, including literature justifications for any proxies used. See Appendix I for units and summary statistics for specific variables.

We encourage future research to move beyond this baseline with finer-scaled or newly collected data. For example, this study could be reproduced at the district level, using the methodology applied here in addition to following Müller and Zeller (2002) and Munroe and Müller (2007). Tree cover loss data and microeconomic data can be aggregated to 1 km-grid cells, and a small micro-survey of households should be conducted if the fine-scaled GSOV household survey data is not acquired. This would also enable more effective use of the Forestry Data Sharing System database.

## Appendix III: Understanding the Moran’s I Outputs

Univariate and bivariate cluster and significance maps are presented in this study. Here we further describe the interpretation of these maps.

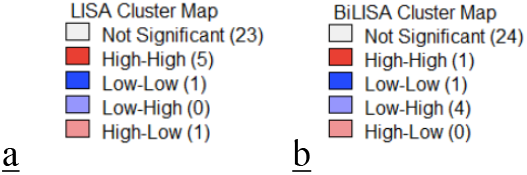

### <u>a.</u> Univariate (Local Indicator of Spatial Association: LISA)

For the univariate statistic, a variable is compared to its own spatial lag. The spatial lag is essentially an average of the values of the variable for the districts surrounding a given district, based on contiguous neighboring districts in this study. The first “High” or “Low” is thus the x-axis of the Moran’s I plot, which is the variable, and the second “High” or “Low” is the y-axis, which is its spatial lag. For example, for the univariate local Moran’s I of precipitation, High-High (dark red) indicates a district of high precipitation surrounded by districts of low precipitation. Low-Low is a district of low precipitation surrounded by districts of low precipitation. Low-High is a district of low precipitation surrounded by districts of high precipitation. Finally, High-Low is a district of high precipitation surrounded by districts of low precipitation. The green significance maps show the level of statistical significance of these clusters. Only plots with an overall local univariate Moran’s I statistic (presented above the x-y plot) significant at a threshold of 5% were included in this study.

### <u>b.</u> Bivariate (Bivariate Local Indicator of Spatial Association: BiLISA)

For the bivariate statistic, a variable is compared to the spatial lag of another variable, which is based on district contiguity (or k nearest neighbor) in this study. Similar to the univariate Moran’s I, the first “High” or “Low” is the x-axis of the Moran’s I plot, which is the non-spatial variable for a given district, and the second “High” or “Low” is the y-axis, but in the bivariate version this is the spatial lag of a different variable. For example, for the bivariate local Moran’s I of irrigated agriculture relative to tree cover loss, High-High (dark red) indicates a district of high irrigated agriculture surrounded by districts of high tree cover loss. Low-Low is a district of low irrigated agriculture surrounded by districts of low tree cover loss. Low-High is a district of low irrigated agriculture surrounded by districts of high tree cover loss. Finally, High-Low is a district of high irrigated agriculture surrounded by districts of low tree cover loss. The nature of the spatial lag is such that the bivariate statistic incorporates the neighborhood of districts but not the specific district itself, and as such must be cautiously interpreted; these statistics, however, are nonetheless useful for understanding the spatial distribution of economic and socioeconomic variables relative to tree cover loss. The green significance maps show the level of statistical significance of these clusters. Only plots with an overall local univariate Moran’s I statistic (presented above the x-y plot) significant at a threshold of 5% were included in this study.

For both map types, numbers in parentheses (e.g., Not Significant (23) or High-High (5)) indicate the number of observations within that category.

See Anselin (1988; 2018), Anselin et al. (2002), Anselin et al. (2008), Lee (2001), and Fotheringham et al. (2000) for further information on these statistics and their methods.

